# The calcium pump PMCA4b promotes epithelial cell polarization and lumen formation

**DOI:** 10.1101/2024.01.20.576436

**Authors:** Sarolta Tóth, Diána Kaszás, János Sónyák, Anna-Mária Tőkés, Rita Padányi, Béla Papp, Réka Nagy, Kinga Vörös, Tamás Csizmadia, Attila Tordai, Ágnes Enyedi

**Author notes:** Corresponding authors: Ágnes Enyedi, Department of Transfusion Medicine, Semmelweis University, H-1089 Budapest, Hungary;, Sarolta Tóth, Department of Transfusion Medicine, Semmelweis University, H-1089 Budapest, Hungary.

## Abstract

Loss of epithelial cell polarity and tissue disorganization are hallmarks of carcinogenesis, in which Ca^2+^ signaling plays a significant role. Here we demonstrate that the plasma membrane Ca^2+^ pump PMCA4 (ATP2B4) is downregulated in luminal breast cancer, and this is associated with shorter relapse-free survival in patients with luminal A and B1 subtype tumors. Using the MCF-7 breast cancer cell model we show that PMCA4 silencing results in the loss of cell polarity while a forced increase in PMCA4b expression induces cell polarization and promotes lumen formation in 2D and 3D cell cultures. We identify Arf6 as a novel regulator of PMCA4b endocytic recycling essential for PMCA4 regulated lumen formation. Silencing of the single *pmca* gene in *Drosophila melanogaster* larval salivary gland destroys lumen morphology suggesting a conserved role of PMCAs in lumen morphogenesis. Our findings point to a novel role of PMCA4 in controlling epithelial cell polarity, and in the maintenance of normal glandular tissue architecture.

## Introduction

Breast cancer is the most commonly occurring cancer in women, representing about 10% of all cancers in the female population worldwide (McGuire et al, 2015). The majority of breast cancers are estrogen (ER+) and progesterone (PgR+) hormone receptor-positive, of which the luminal A (LUMA) subtype is the least aggressive (low grade) while the more proliferative (Ki67-high) luminal B1 (LUMB1) and human epidermal growth factor receptor 2-positive (HER2+) luminal B2 (LUMB2) subtypes are of higher grades.

Intracellular Ca^2+^ homeostasis is frequently altered in cancer cells, and this can play a role in tumor initiation, progression, angiogenesis and metastasis (Cui et al., 2017; Monteith et al., 2017). Plasma membrane calcium ATPases (PMCA/ATP2B) regulate intracellular Ca^2+^ concentration by pumping Ca^2+^ out of the cytoplasm into the extracellular space in order to maintain the exceedingly low cytosolic Ca^2+^ concentration. In mammalian cells four genes (*ATP2B1-4*) encode PMCA1-4 proteins, and as a result of alternative splicing, more than 20 PMCA isoforms have been described. PMCA1 is the house-keeping form that together with PMCA4 is expressed in all cell types while PMCA2 and PMCA3 expressions are tissue specific (Krebs, 2015; Padanyi et al., 2016). Both PMCA2 and PMCA4 play a role in the adult mammary gland, but their functions are quite different (Reinhardt et al., 2000). PMCA2 is highly expressed during the lactating period and is responsible for setting Ca^2+^ concentration in the milk (Reinhardt et al., 2004). In contrast, PMCA4 is upregulated before lactation and during mammary gland involution (Reinhardt et al., 2000; Reinhardt and Lippolis, 2009). It has been well established that serum estradiol levels gradually increase during pregnancy, and we showed that the “b” splice variant of PMCA4 (PMCA4b) expression was highly dependent on estradiol concentrations in ER-α positive MCF-7 breast cancer cells, suggesting a role for this pump in mammary gland development (Varga et al., 2018).

Alterations in PMCA expressions have been reported in various cancer types (Islam, 2020; Monteith et al., 2017). Among these, PMCA4b was shown to be down-regulated in ER^+^ luminal-type breast cancer cells whereas its expression was relatively high in basal-type cells with, however, a predominant localization to intracellular compartments (Varga et al., 2018). Low expression of PMCA4b was detected in highly metastatic BRAF-mutant melanoma cell lines, in which re-expression of PMCA4b was associated with decreased migration and metastatic activities suggesting a metastasis suppressor role for this PMCA variant (Hegedus et al., 2017a).

Luminal epithelial cells display apical-basal polarity that is regulated by specific protein modules under the control of the Par, Crumbs and Scribble polarity protein complexes (Martin et al., 2021). Lumen formation is an essential step in mammary gland development, and loss of lumens is a hallmark of tumorigenesis (Halaoui et al., 2017). It has been shown that primary mammary epithelial cells isolated from mice were able to form lumen-containing acini on reconstituted basement membrane matrix, however, after oncogene induction, the cells lost their epithelial polarity and lumen-forming capability (Jechlinger et al., 2009). In polarized cells directed vesicular transport is essential to ensure proper protein and lipid composition of specialized membrane compartments (Jewett and Prekeris, 2018). The ADP-ribosylation factor 6 (Arf6) small GTPase protein regulates membrane trafficking pathways (Li et al., 2017), and it has been implicated in cell polarization (Ikenouchi and Umeda, 2010; Osmani et al., 2010; Shteyn et al., 2011) and lumenogenesis (Clancy et al., 2018; Tushir et al., 2010). Although Arf6 is essential for cell polarization, Arf6 hyperactivity can promote invasiveness and metastatic activity of cancer cells (Li et al., 2017; Nikolatou et al., 2023) by increasing their motility (Zobel et al., 2018).

In the present study we show that PMCA4(b) (assessed by a PMCA4-specific antibody) is downregulated in luminal-type breast carcinoma tissue samples, and that the expression level of the *ATP2B4* gene correlates with patient survival. We demonstrate that PMCA4b promotes polarization and lumen formation of ER^+^ MCF-7 breast cancer cells, and this requires the previously identified di-leucine endocytic motif at the C-terminus of the pump (Antalffy et al., 2013). Here we find that PMCA4b shows strong colocalization with wild-type and constitutively active forms of Arf6, moreover inhibition of Arf6 leads to the intracellular accumulation of PMCA4b that defines Arf6 as a new regulator of PMCA4b trafficking. To investigate the evolutionarily conserved role of PMCAs in lumen formation we silenced the single *Drosophila melanogaster* PMCA (dmPMCA) in larval salivary gland, and this disrupted lumen morphology and induced a severe secretion defect suggesting a conserved role of PMCAs in lumen formation and secretion.

## Results

### PMCA4 is downregulated in HR^+^ breast cancer

Besides an analysis of publicly available gene expression datasets (Varga et al., 2018), no clinically relevant data have been published on PMCA4 expression in breast cancer. Since a majority of all breast cancer cases (about 70-80%) are hormone receptor-positive (HR^+^) (Harbeck et al., 2019), we investigated PMCA4 protein abundance in HR^+^ breast carcinoma tissue samples (Suppl. Table 1) using the PMCA4-specific antibody JA9 (JA9 does not discriminate between PMCA4 “a” and “b” splice variants) (Caride et al., 1996), and compared the results to those of normal breast tissue samples obtained after breast reduction surgery. We found that PMCA4 was highly expressed in the ductal epithelium of normal breast tissue and in more differentiated regions of tumor samples when compared to fully de-differentiated regions of the tumor (Fig. 1A). In 76% of the breast cancer cases (83 out of 109 tissue samples) less than 5% of the tumor cells showed PMCA4 staining. Analyzing separately the HR^+^ subtypes, no statistically significant differences were found between the LUMA and LUMB cases (Fig. 1B, Suppl. Table 1). Data from the Clinical Proteomic Tumor Analysis Consortium (Chandrashekar et al., 2022) (CPTAC) (https://ualcan.path.uab.edu) confirmed these findings and showed significantly reduced PMCA4 protein levels in luminal breast cancer types compared to normal breast tissue. In contrast to PMCA4, PMCA1 expression was not different in the same comparison (Fig. 1C).

**Figure 1.**
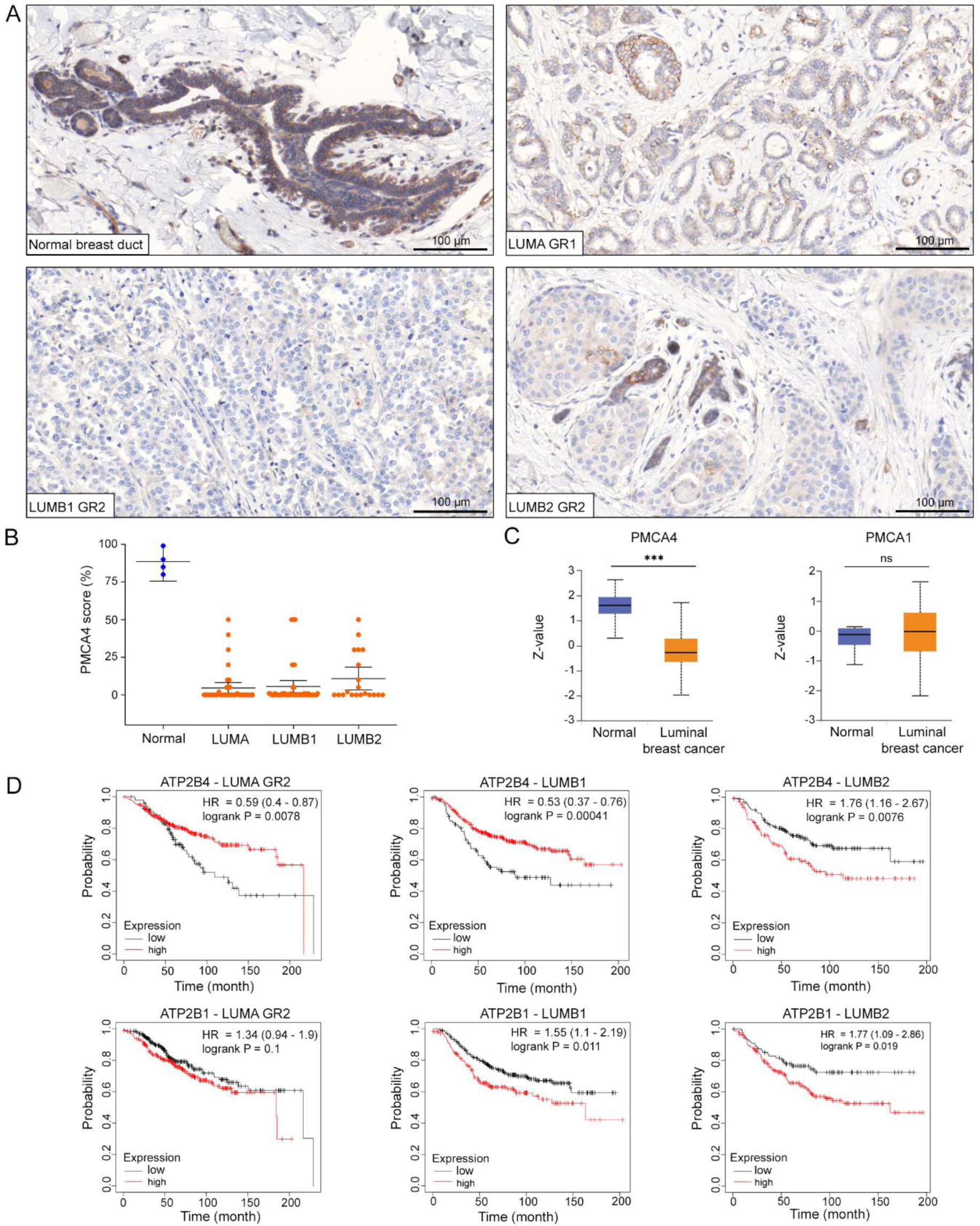
PMCA4 is downregulated in HR^+^ breast cancer subtypes. **A** PMCA4-specific antibody staining of normal and neoplastic LUMA, LUMB1 and LUMB2 subtype mammary gland tissue samples. GR1 or 2 indicates histological grade 1 or grade 2. **B** Fraction of cancer cells showing positive membranous PMCA4 staining; n_(normal)_=4 (obtained after breast reduction surgery), n_(LUMA)_=39, n_(LUMB1)_=50, n_(LUMB2)_=20. **C** Protein expression of PMCA4 (NP_001675.3:S328) and PMCA1 (NP_001673.2:S1182) in normal breast tissue and luminal breast cancer subtypes. Data derived from the Clinical Proteomic Tumor Analysis Consortium (CPTAC) (http://ualcan.path.uab.edu) that are expressed as z-values and analyzed by Student’s t test; n_(PMCA4b/normal)_=18, n_(PMCA4b/luminal)_=64, n_(PMCA1b/normal)_=18, n_(PMCA4b/luminal)_=64; ***p<0.001, ^ns^p>0.05; ns indicates non-significant difference. **D** Kaplan–Meier relapse free survival analysis in breast cancer patients with LUMA, LUMB1 or LUMB2 subtype tumors with low and high *ATP2B4*/PMCA4 and *ATP2B1*/PMCA1 expression levels. GR 2 indicates histological grade 2. Data were collected by the online survival analysis tool (http://www.kmplot.com) using microarray data analysis; n_(ATP2B4/LUMA-GR2/low)_=97, n_(ATP2B4/LUMA-GR2/high)_=297, n_(ATP2B4/LUMB1/low)_=99, n_(ATP2B4/LUMB1/high)_=301, n_(ATP2B4/LUMB2/low)_=150, n_(ATP2B4/LUMB2/high)_=108, n_(ATP2B1/LUMA-GR2/low)_=257, n_(ATP2B1/LUMA GR2/high)_=254, n_(ATP2B1/LUMB1/low)_=239, n_(ATP2B1/LUMB1/high)_=161, n_(ATP2B1/LUMB2/low)_=89, n_(ATP2B1/LUMB2/high)_=169.

To assess the prognostic significance of ATP2B4 (PMCA4) expression in the clinical outcome of HR^+^ breast carcinoma subtypes, the publicly available KM Plotter Online Tool (Gyorffy, 2021) (www.kmplot.com) was used. According to this database, high ATP2B4 gene expression was associated with greater probability of relapse-free survival (RFS) in grade 2 LUMA and in LUMB1 tumors, whereas an inverse correlation was observed in the HER2-expressing LUMB2 subtype, in which poorer outcomes were associated with higher ATP2B4 expression. It is worth noting that in grade 1 and 3 LUMA tumors the effects were not significant (Suppl. Fig. 1). In contrast to ATP2B4, no statistically significant association was detected between low and high ATP2B1 gene expression and RFS in the LUMA and LUMB1 breast carcinoma subtypes, while in the LUMB2 subtype high ATP2B1 expression (PMCA1) level was associated with a lower probability of RFS as also seen for high ATP2B4 (PMCA4) expression (Fig. 1D).

### PMCA4b is needed for the plasma membrane localization of E-cadherin in MCF-7 breast cancer cells

High PMCA4 protein levels in the normal mammary gland and its nearly complete loss in luminal subtype tumors suggest that PMCA4 is involved in the development and/or maintenance of the normal mammary tissue. The PMCA4b splice variant is an ubiquitous form of PMCA4, and this isoform has been found in several luminal-type breast cancer cells including the estrogen-sensitive MCF-7 cell line (Varga et al., 2018; Varga et al., 2014). Although these cells express low levels of PMCA4b, they retain most of their epithelial features since they express the epithelial marker E-cadherin but not the mesenchymal markers N-cadherin and vimentin (Comsa et al., 2015). To understand the role of PMCA4(b) loss during tumor progression, the endogenous PMCA4b was silenced by a PMCA4-specific short hairpin RNA (sh-RNA), or the GFP-tagged PMCA4b and its endocytosis-defective mutant form PMCA4b^LA^ (Antalffy et al., 2013) were stably over-expressed in MCF-7 cells. We found that neither silencing nor over-expression of PMCA4b affected the expression levels of the epithelial and mesenchymal markers E-cadherin, N-cadherin and vimentin (Fig. 2A, Suppl. Fig. 2A), respectively, or influenced cell growth (Suppl. Fig. 2B). However, silencing PMCA4 induced internalization of E-cadherin, resulting in its reduced plasma membrane localization. In contrast, the parental, GFP-PMCA4b- or the trafficking-mutant GFP-PMCA4b^LA^-expressing MCF-7 cells displayed more prominent plasma membrane localization of E-cadherin (Fig. 2B-E, Suppl. Fig. 2C). This suggests that loss of PMCA4 may induce partial epithelial-mesenchymal transition (EMT) through E-cadherin internalization without affecting the overall expression of the EMT markers.

**Figure 2.**
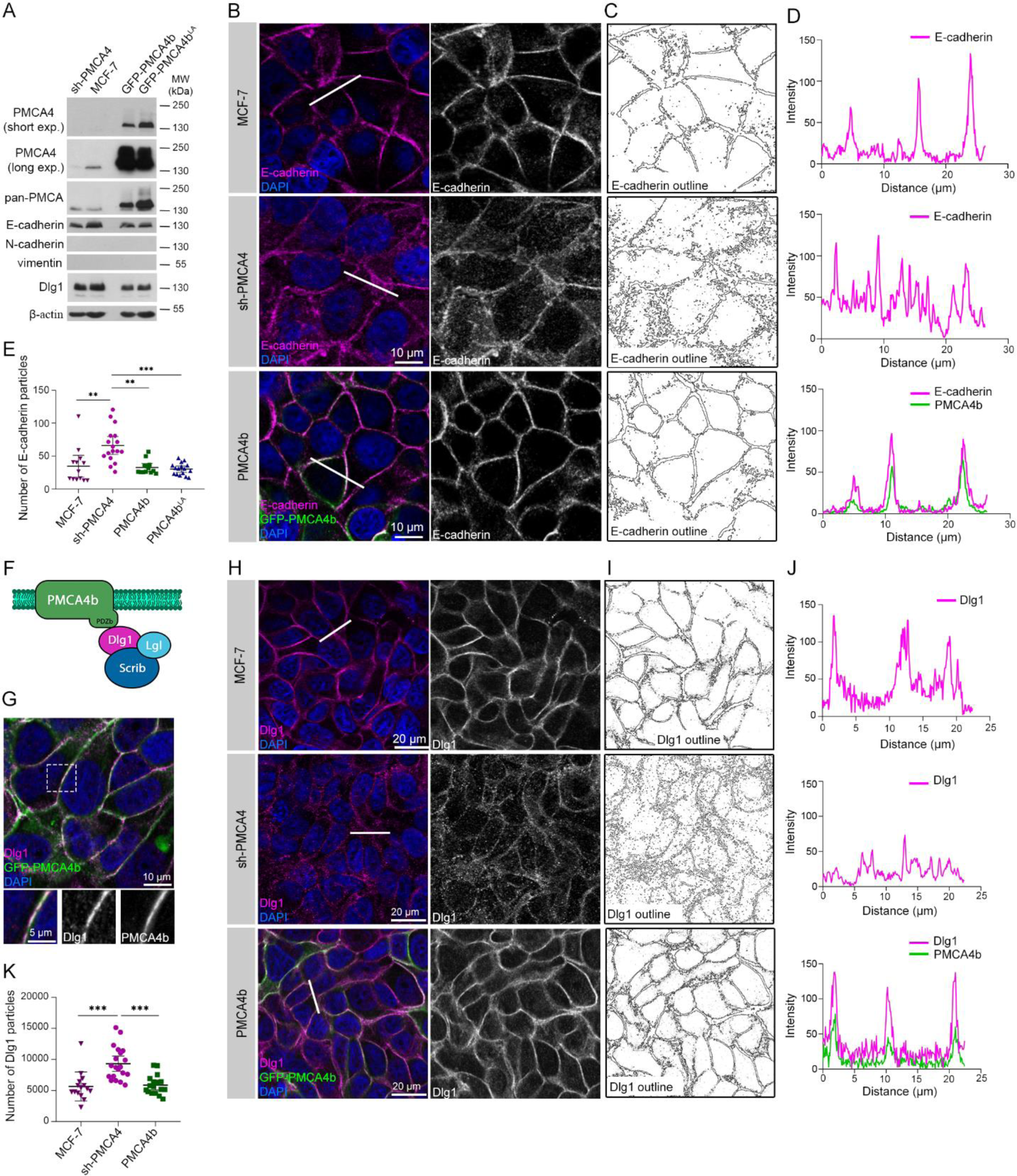
PMCA4b is necessary for the plasma membrane localization of E-cadherin and Dlg1**. A** Western blot analysis of parental, PMCA4 specific shRNA (sh-PMCA4), GFP-PMCA4b and GFP-PMCA4b^LA^-expressing MCF-7 cell lines; exp. denotes time of exposure. **B** E-cadherin immunostaining of parental, PMCA4 specific shRNA (sh-PMCA4) and GFP-PMCA4b-expressing MCF-7 cell lines. DAPI (4′, 6-diamidino-2-phenylindole, blue) labels nuclei. **C** Outline of the E-cadherin positive cell compartments on B generated by the Image J software. **D** Fluorescence intensity profiles of E-cadherin and GFP-PMCA4b corresponding to the white line shown in B. **E** Statistical analysis of the number of E-cadherin positive particles in parental, PMCA4 specific shRNA (sh-PMCA4), GFP-PMCA4b and GFP-PMCA4b^LA^-expressing MCF-7 cell lines. Graph displays means with 95% confidence intervals (CI); data were collected from 2 independent experiments, n_(MCF-7)_=13, n_(sh-PMCA4)_=17, n_(PMCA4b)_=13, n_(PMCA4b_^LA^_)_=16. Data were analyzed with Kruskal-Wallis and Dunn’s multiple comparisons tests, adjusted p values: **p<0.01, ***p<0.001. Non-significant differences are not indicated. **F** Model of the interaction between PMCA4b and the Scrib-Dlg1-Lgl polarity complex. PDZb denotes the PDZ-binding sequence motif of PMCA4b. **G** Dlg1 immunostaining of GFP-PMCA4b-expressing MCF-7 cells. Insets show magnification of the area framed by the dotted line. DAPI labels nuclei. **H** Dlg1 immunostaining of parental, PMCA4 specific shRNA (sh-PMCA4) and GFP-PMCA4b-expressing MCF-7 cell lines. DAPI labels nuclei. **I** Outline of the Dlg1 positive cell compartments on H generated by the ImageJ software. **J** Fluorescence intensity profiles of Dlg1 and GFP-PMCA4b corresponding to the white line shown on panel H. **K** Statistical analysis of the number of Dlg1 positive particles in parental, PMCA4-specific shRNA (sh-PMCA4) and GFP-PMCA4b-expressing MCF-7 cell lines. Graph displays means with 95% CI; data were collected from 2 independent experiments, n_(MCF-7)_=15, n_(sh-PMCA4)_=20, n_(PMCA4b)_=17; data were analyzed with Kruskal-Wallis and Dunn’s multiple comparisons tests; adjusted p values: ^ns^p>0.05, ***p<0.001; non-significant difference is not labeled.

### PMCA4b regulates epithelial cell polarity

E-cadherin is involved in cell-cell adhesion and cell polarization partly through its interaction with scribble (Scrib), a member of the apico-basal Scribble polarity complex that consists of lethal giant larvae (Lgl), discs large 1 (Dlg1) and Scrib proteins (Wang et al., 2018). It has been known for a long time that the C-terminal PDZ-binding sequence motif ETSV of PMCA4b binds to the PDZ-domains 1 and 2 of the scaffold protein Dlg1 (DeMarco and Strehler, 2001) (Fig. 2F), and this way it can participate in cell polarization. Here we show that Dlg1 is highly expressed in MCF-7 cells and its expression is not affected by either PMCA4b silencing or over-expression (Fig. 2A). Fluorescence immunostaining of PMCA4b-expressing MCF-7 cells show co-localization between PMCA4b and Dlg1 at the plasma membrane, as expected (Fig. 2G), while PMCA4-silencing is associated with preferential cytoplasmic Dlg1-localization (Fig. 2H-K), suggesting that PMCA4b is involved in targeting Dlg1 to the plasma membrane.

Ezrin is an F-actin-binding protein that is known to localize to the apical plasma membrane of polarized epithelial cells (Casaletto et al., 2011). The overall high level and/or atypical localization of ezrin in tumor cells were shown to correlate with poor prognosis in breast cancer (Jeong et al., 2019; Sarrio et al., 2006). While expression of sh-PMCA4, PMCA4b or the trafficking mutant PMCA4b^LA^ in MCF-7 cells did not alter ezrin protein levels (Fig. 3A), variations in PMCA4b expression showed strikingly different localization patterns. In contrast to the PMCA4-silenced or trafficking mutant PMCA4b^LA^-expressing MCF-7 cells where a substantial proportion of ezrin was localized to intracellular compartments, PMCA4b over-expression resulted in a significantly more prominent asymmetric ezrin localization when compared to the parental MCF-7 cells (Fig. 3B-D). These data suggest that PMCA4b abundance affects cell polarization and show that not only PMCA4b expression but also its proper trafficking is required to accomplish cell polarity.

**Figure 3.**
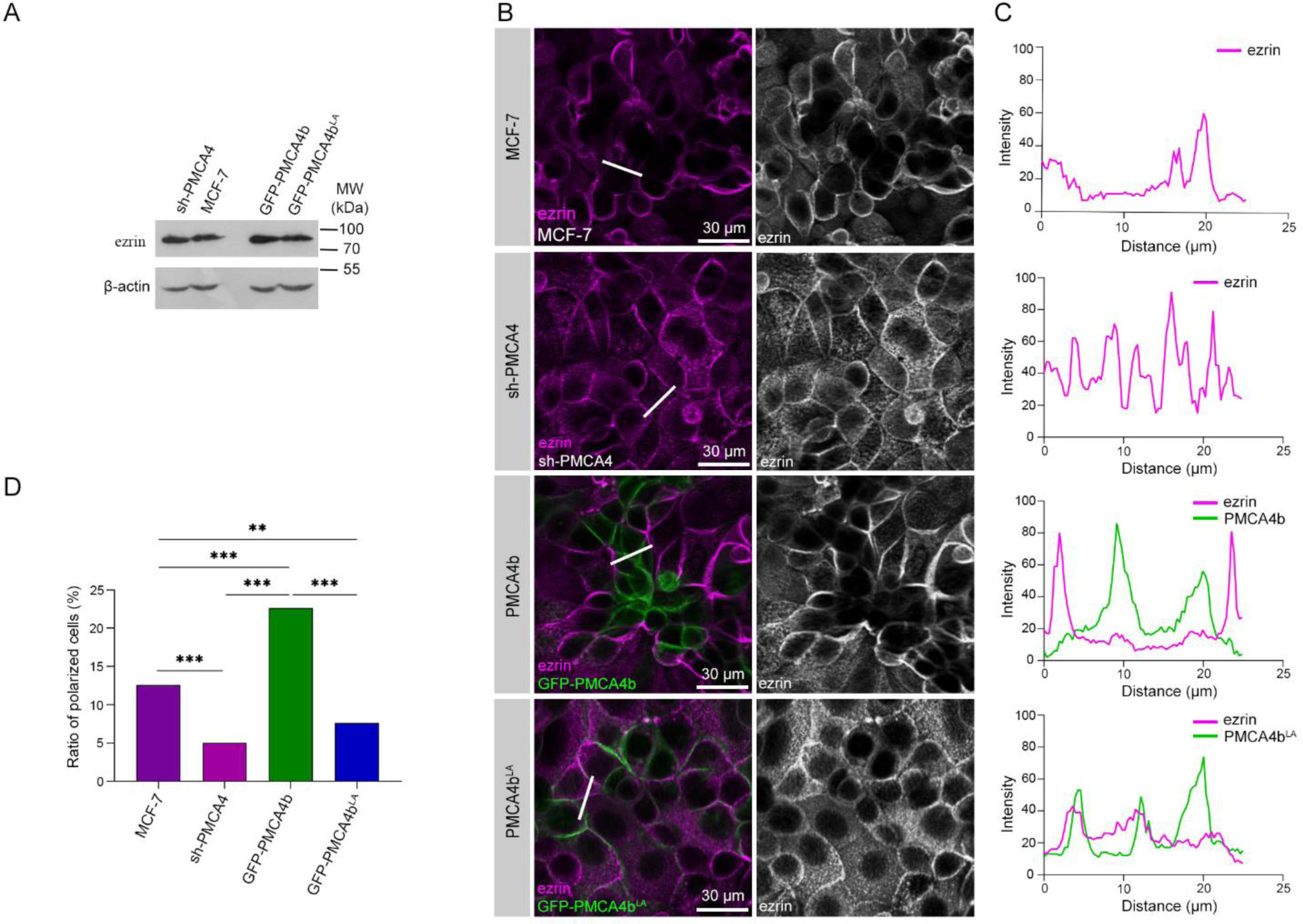
PMCA4b induces polarized ezrin distribution. **A** Western blot analysis for ezrin in parental, PMCA4 specific shRNA (sh-PMCA4), GFP-PMCA4b and GFP-PMCA4b^LA^-expressing MCF-7 cell lines. **B** Ezrin immunostaining of parental, PMCA4-specific shRNA (sh-PMCA4), GFP-PMCA4b and GFP-PMCA4b^LA^-expressing MCF-7 cell lines. DAPI labels nuclei. **C** Fluorescence intensity profiles of ezrin and GFP-PMCA4b corresponding to the white lines shown on panel B. **D** Statistical analysis of cell polarization rate based on ezrin distribution in parental, PMCA4-specific shRNA (sh-PMCA4), GFP-PMCA4b and GFP-PMCA4b^LA^-expressing MCF-7 cell cultures. Data were collected from 16 fields of view from 2 independent experiments; n_(MCF-7)_=1589, n_(sh-PMCA4)_=1699, n_(PMCA4b)_=1462, n_(PMCA4b_^LA^_)_=1511. Data were analyzed with Chi-square test, p values: **p<0.01, ***p<0.001; non-significant difference is not labeled.

### PMCA4b enhances polarized vesicular trafficking

Epithelial cells require polarized vesicular trafficking for the proper distribution of proteins and lipids at specific plasma membrane domains to establish and maintain apico-basal cell polarity (Banushi et al., 2023). Wheat germ agglutinin (WGA) has been widely used to follow endosomal trafficking in live cells. Upon binding to N-acetylglucosamine and sialic acid moieties on the extracellular side of plasma membrane proteins WGA can be internalized and transported through the endosomal pathways (Liu et al., 2011). We performed a WGA uptake assay and found that PMCA4b-expressing cells collected WGA-positive vesicles to one side of the plasma membrane (Video 1) as a sign of polarized vesicular trafficking while these vesicles were located randomly in the cytoplasm of the parental, the sh-PMCA4- and the PMCA4b^LA^-expressing cells (Fig. 4A-D, Video 2-3, Suppl. Fig. 4). Furthermore, PMCA4b strongly co-localized with WGA-positive vesicles in contrast to the PMCA4b^LA^ mutant that clearly separated from the WGA-positive puncta (Fig. 4E) confirming the importance of PMCA4b trafficking in cell polarization.

**Figure 4.**
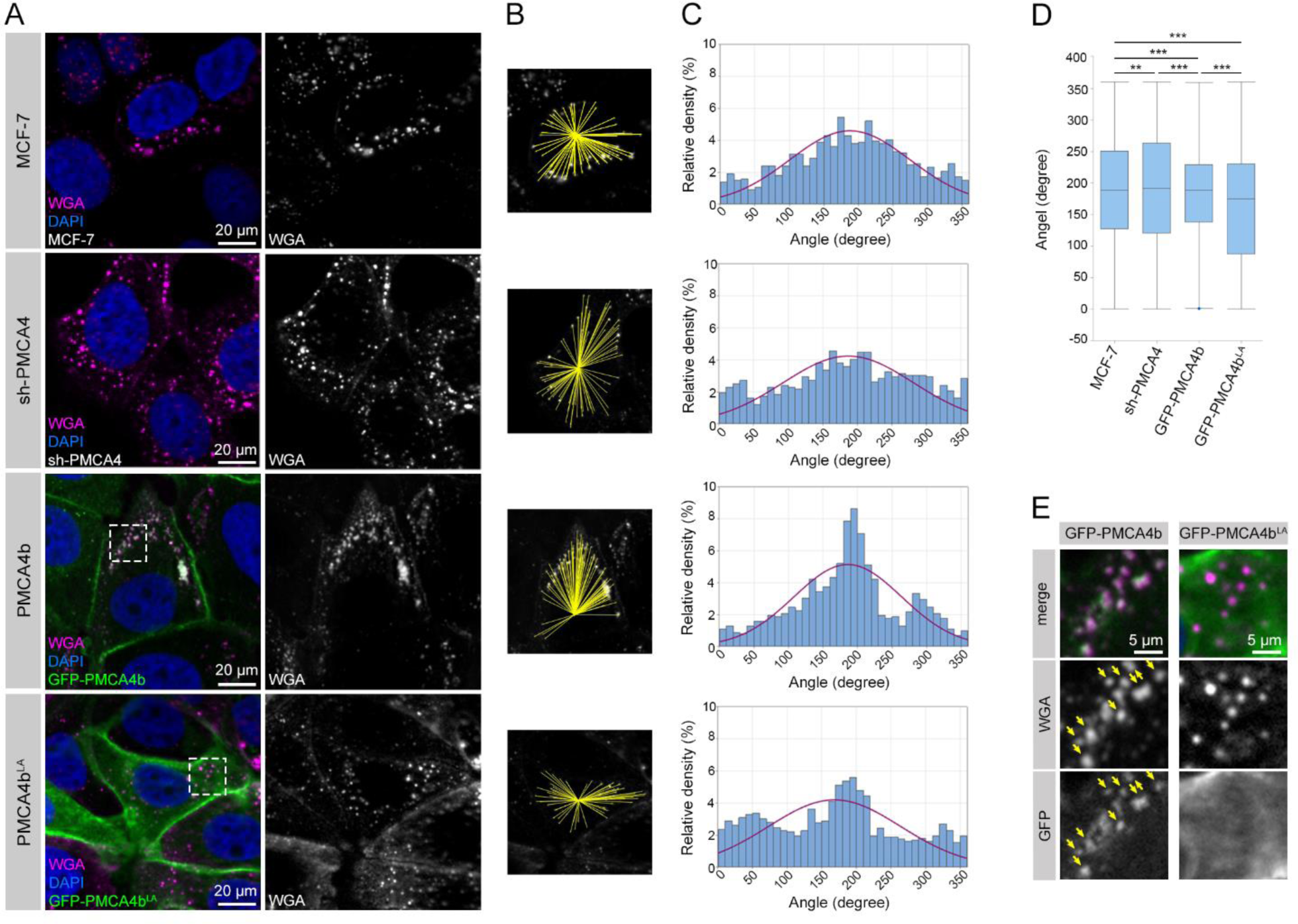
PMCA4b promotes polarized vesicular trafficking. **A** WGA uptake assay was performed on parental, PMCA4-specific shRNA (sh-PMCA4), GFP-PMCA4b and GFP-PMCA4b^LA^-expressing MCF-7 cell lines. After 10 minutes incubation with WGA, cells were washed and transferred to a 37 °C incubator for 20 minutes. Dashed white squares indicate areas for magnification on D. DAPI labels nuclei. **B** Yellow lines connect the center of the cell nucleus with WGA positive endocytic vesicles accumulated in the cytosol. **C** Intracellular distribution analysis of WGA positive vesicles. Red line indicates the distribution fit curve. Data were collected from 2 independent experiments, *n_(MCF-7)_*=1214, n_(sh-PMCA4)_=922, n_(PMCA4b)_=1021, n_(PMCA4b_^LA^_)_=1493 from 15-28 cells, and analyzed as described in the Methods section. **D** Statistical analysis of the intracellular distribution of WGA vesicles and illustrated by a box plot diagram. The same data as in C were analyzed with Levene’s test. Data were the same as C and analyzed with Levene’s test. P values: **P<0.01, ***P <0.001; non-significant difference is not labeled. **E** Magnification of the areas indicated by the dashed white squares on panel A. Yellow arrows indicate colocalized WGA and PMCA4b positive vesicles.

### Arf6 regulates intracellular trafficking of PMCA4b

PMCAs require complex formation with neuroplastin or CD147/basigin for their proper plasma membrane localization and function (Korthals et al., 2017; Schmidt et al., 2017). It has been demonstrated that internalization and recycling of CD147 is mediated by Arf6 (Qi et al., 2019), and Arf6 has been implicated in cell polarization (Ikenouchi and Umeda, 2010). To study the role of Arf6 in PMCA4b trafficking we transiently over-expressed wildtype Arf6 and its constitutively active mutant form Arf6^Q67L^ in PMCA4b-MCF-7 and PMCA4b-HEK293 cells. Strong co-localization between PMCA4b and wild type Arf6 was seen near the plasma membrane and intracellular compartments (Fig. 5A) while the mutant Arf6^Q67L^ caused intracellular accumulation of PMCA4b in both cell types (Fig. 5B), similarly to that described for other known Arf6 cargo proteins (Porat-Shliom et al., 2013; Sannerud et al., 2011). To inhibit Arf6 we treated PMCA4b-expressing MCF-7 cells with NAV2729 (Yoo et al., 2016). After 24 hours of the treatment, we observed an enrichment of PMCA4b-CD147 positive vesicles near the plasma membrane, and after 48 hours PMCA4b-CD147 vesicles accumulated in the cytoplasm (Fig. 5C) suggesting that Arf6 was involved in the endocytic recycling of the PMCA4b-CD147 complex. These results indicate that Arf6 is a novel regulator of PMCA4b-CD147 endosomal trafficking (Fig. 5D).

**Figure 5.**
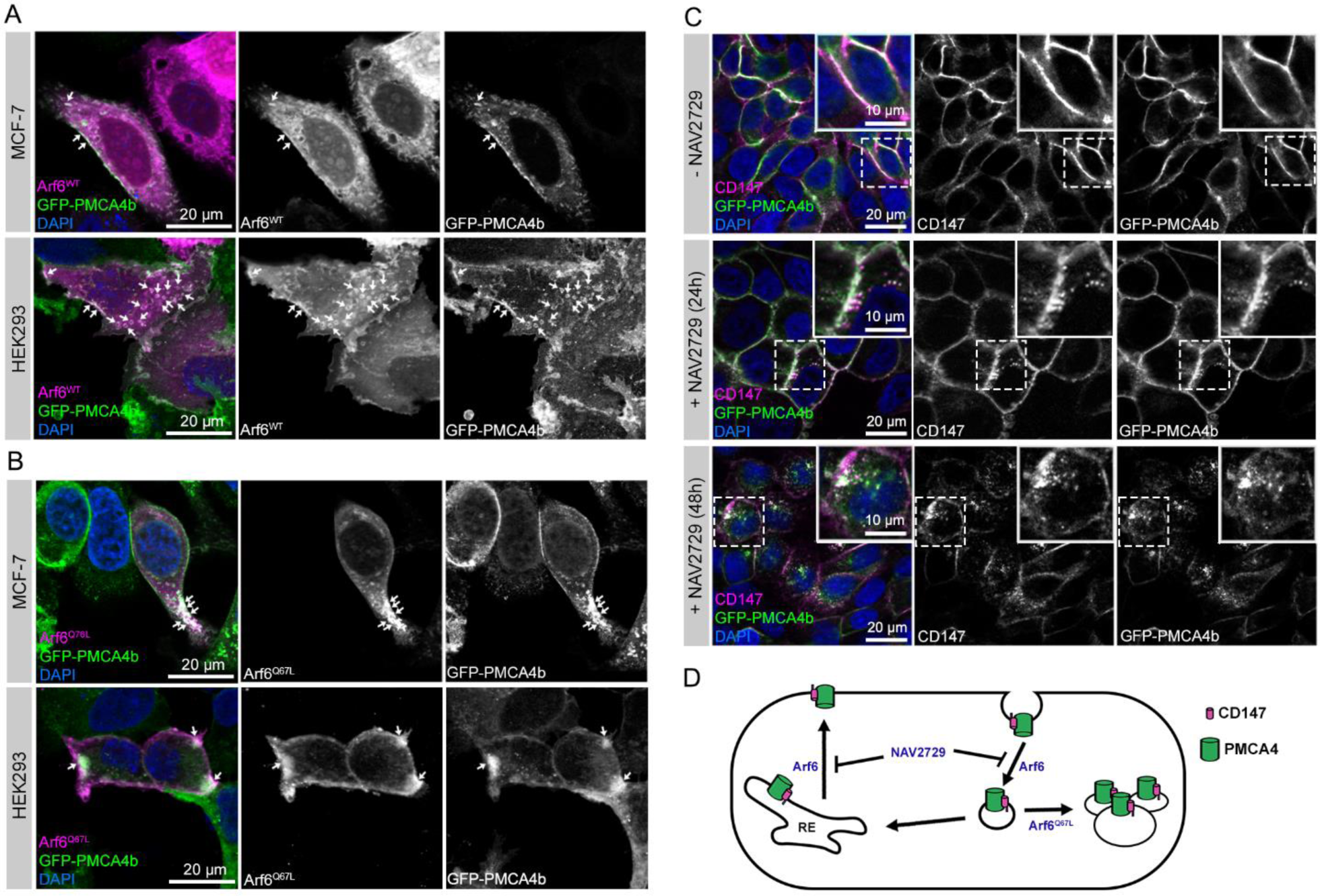
Arf6 regulates endocytic recycling of PMCA4b. **A-B** Transfection of GFP-PMCA4b-expressing MCF-7 and HEK293 cell lines with wild-type (WT) and Q67L mutant form of Arf6. White arrows point to colocalizing areas. **C** CD147 immunostaining of non-treated and 24 or 48 hours NAV2729-treated GFP-PMCA4b-expressing MCF-7 cells. Insets show magnified areas indicated by the dashed white rectangles. **D** A scheme depicting the role of Arf6 in PMCA4b trafficking. RE: recycling endosome.

### PMCA4b promotes pre-lumen formation in 2D cultures of MCF-7 cells

The establishment and maintenance of epithelial cell polarity are hallmarks of healthy mammary tissue organization featured by a complex ductal network. However, during neoplastic transformation the polarity of neoplastic mammary epithelial cells is lost and their capability to form mature mammary ducts with well-defined central lumens is also compromised (Chatterjee and McCaffrey, 2014). We found that in highly confluent 2D cultures of MCF-7 cells PMCA4b over-expression led to the formation of significantly more pre-lumen-like structures resembling cellular intermediates of lumen formation than in the case of the parental, the sh-PMCA4 or the trafficking mutant PMCA4b^LA^-expressing cells (Fig. 6A, D, E). PMCA4b colocalized with phalloidin labeled actin positive vesicles and WGA positive endosomes near pre-lumens (Fig. 6B, C) indicating that PMCA4b participates in the dynamic vesicular trafficking characteristic of *de novo* lumen formation (Datta et al., 2011; Levic and Bagnat, 2023).

**Figure 6.**
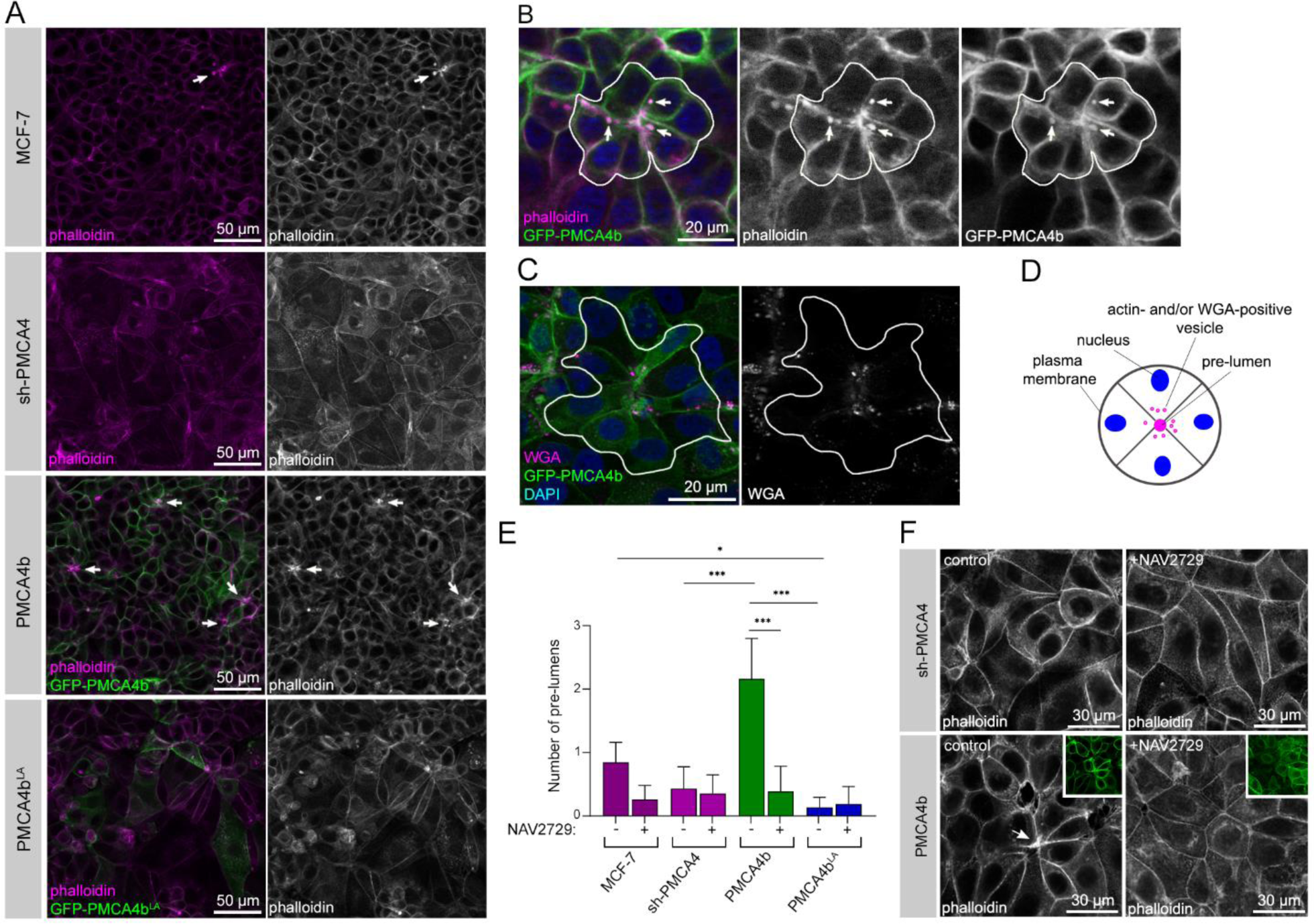
PMCA4b promotes pre-lumen formation in 2D cultures of MCF-7 cells. **A** Phalloidin staining of parental, PMCA4 specific shRNA (sh-PMCA4), GFP-PMCA4b and GFP-PMCA4b^LA^-expressing MCF-7 cell lines. White arrows point to the pre-lumens. **B** Phalloidin staining of GFP-PMCA4b-expressing MCF-7 cells. The white line encircles a pre-lumen. White arrows point to phalloidin and PMCA4b double-positive vesicles. **C** WGA uptake assay on GFP-PMCA4b-expressing MCF-7 cells with 20 minutes incubation after WGA removal. Pre-lumen creating cells are encircled with white line. **D** A schematic figure of a typical pre-lumen structure in a 2D cell culture. **E** Statistical analysis of pre-lumen formation in control and NAV2729 treated parental, PMCA4 specific shRNA (sh-PMCA4), GFP-PMCA4b and GFP-PMCA4b^LA^-expressing MCF-7 cell lines. Graph displays means with 95% CI corrected with the total area of the images; data were collected from 2 independent experiments, n_(sh-PMCA4)_=23, n_(sh-PMCA4+NAV2729)_=21, n_(MCF-7)_=26, n_(MCF-7+NAV2729)_=22, n_(PMCA4b)_=25, n_(PMCA4b+NAV2729)_=23, n_(PMCA4b_^LA^_)_=23, n_(PMCA4b_^LA^+_NAV2729)_=17. Data were analyzed with Kruskal-Wallis and Dunn’s multiple comparisons tests, adjusted p values: ^ns^p≥0.05, *p<0.05, **p<0.01 ***p<0.001; non-significant differences are not labeled. **F** Phalloidin staining of control and NAV2729 treated PMCA4 specific shRNA (sh-PMCA4) and GFP-PMCA4b-expressing MCF-7 cell lines. Scaled-down insets show the GFP-PMCA4b signal. The yellow arrowhead points to a pre-lumen.

We treated the PMCA4b-expressing MCF-7 cells with the Arf6 inhibitor NAV2729 and found significantly fewer F-actin labeled pre-lumen structures compared to the non-treated PMCA4b-expressing cells suggesting that Arf6 activity is needed for the positive effect of PMCA4b on pre-lumen formation (Fig. 6E, F, Suppl. Fig. 3).

### PMCA4b promotes central lumen formation in MCF-7 mammospheres

To confirm the role of PMCA4b in lumen formation, we cultured MCF-7 cells in Matrigel for 10 days to create mammospheres and observed that the PMCA4b-expressing cells formed well-defined, F-actin-bordered central lumen-bearing spheroids with a single layer of cells around the lumen (Fig. 6A-C, Suppl. Fig. 4C) similar to that seen in cross-sections of healthy mammary duct (Hassiotou and Geddes, 2013). While the parental MCF-7 cells also formed some central lumen-bearing mammospheres, the lumen structures were not well-defined, and their number was significantly lower compared to the cell cultures with elevated PMCA4b expression (Fig. 6A-C, Suppl. Fig. 4A). Mammospheres generated by sh-PMCA4 or PMCA4b^LA^-expressing MCF-7 cells did not form central lumens; rather, they displayed several small F-actin-rich areas (Fig. 6A-C, Suppl. Fig. 4B, D) resembling the microlumens commonly seen in ductal carcinoma *in situ* (DCIS) (Winchester et al., 2000). These data indicate that both PMCA4b abundance and proper localization was required for appropriately positioned central lumen formation.

Electron microscopy revealed enrichment of secretory vesicles in proximity to the plasma membrane. However, in contrast to the parental and PMCA4b-expressing MCF-7 cells, secretory vesicles - especially the dense core-types - showed excessive accumulation in sh-PMCA4-expressing MCF-7 cells, indicating a pronounced secretion defect in the absence of PMCA4b (Fig. 6D). Interestingly, in PMCA4b^LA^-expressing cells small endocytic vesicles accumulated near the plasma membrane (Fig. 6D) underlining the importance of endosomal trafficking in secretion.

### Drosophila PMCA regulates normal lumen morphology in larval salivary gland

The only *Drosophila melanogaster* (dm)PMCA (UniProt: Q9V4C7) shows 62.7% identity in amino acid sequence with the human PMCA4b protein (UniProt: P23634-6) (Suppl. Fig. 5A). For testing the expression and localization of dmPMCA in the larval salivary gland we used the *pan*-PMCA antibody 5F10 and showed that it was suitable for the detection of dmPMCA because of its highly conserved epitope (Fig. 8 A, C, Suppl. Fig. 5A). Endogenous dmPMCA localized to the basolateral and apical plasma membrane of the salivary gland cells (Fig. 7A) and showed strong co-localization with the *Drosophila melanogaster* Dlg protein (Fig. 7B) in good agreement with the potential PDZ-binding sequence present at the C-terminus of six out of the seven dmPMCA isoforms (Suppl. Fig. 7B).

**Figure 7.**
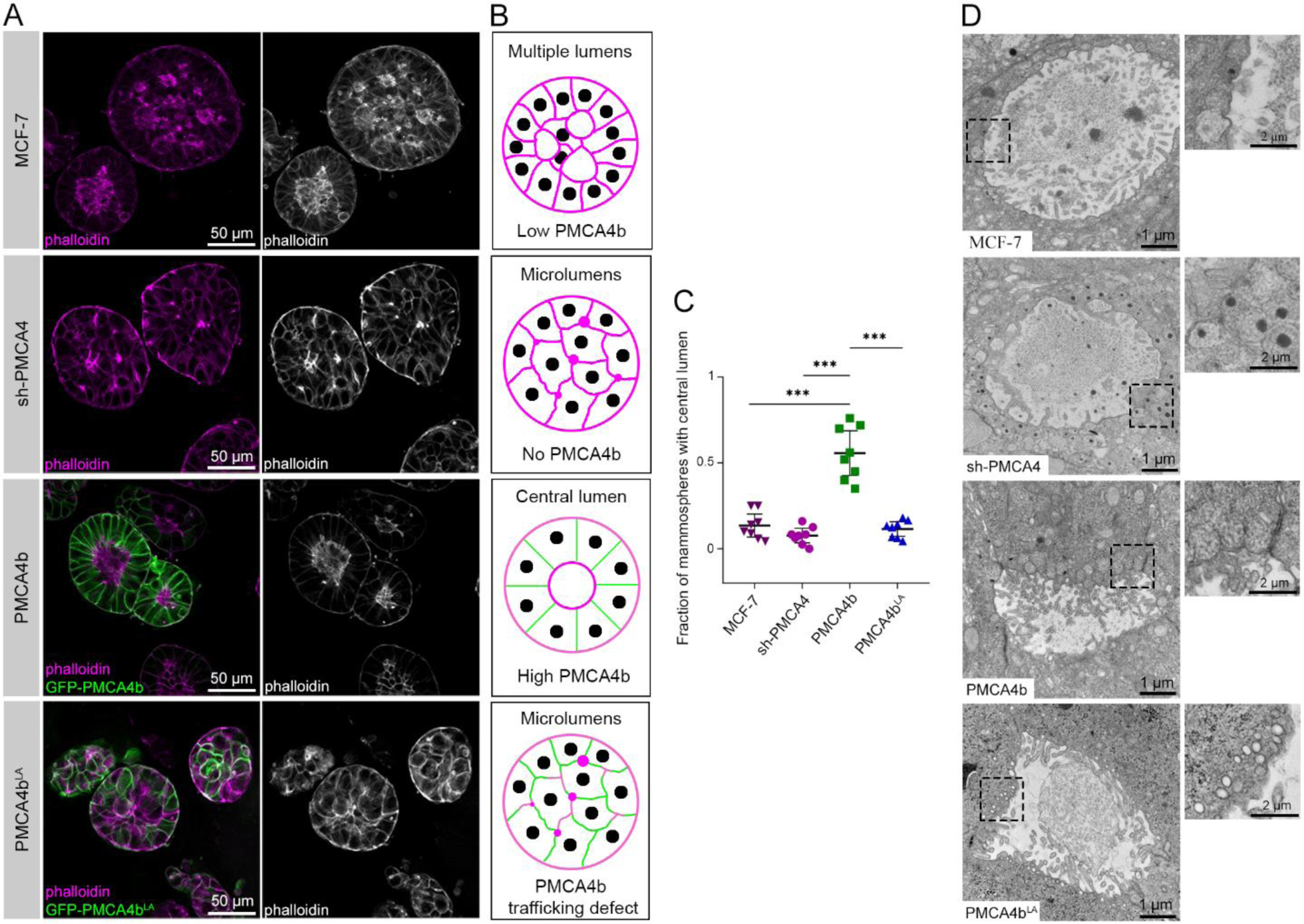
PMCA4b promotes central lumen formation in 3D cultures of MCF-7 cells **A** Phalloidin staining of mammospheres formed by parental, PMCA4-specific shRNA (sh-PMCA4), GFP-PMCA4b and GFP-PMCA4b^LA^-expressing MCF-7 cell lines cultured in Matrigel for 10 days. **B** Schematic figures demonstrating structures of mammospheres based on results on A. **C** Statistical comparisons of the ratios of mammospheres with central lumen in 3D cultures between parental, PMCA4-specific shRNA (sh-PMCA4), GFP-PMCA4b- and GFP-PMCA4b^LA^-expressing MCF-7 cell lines cultured in Matrigel. Graph displays means with 95% CI, *n*=8 images with 10-15 mammospheres per image for each cell type. Data were analyzed with ordinary one-way ANOVA and Tukey’s multiple comparisons tests, adjusted p values: ^ns^p≥0.05, *p<0.05, **p<0.01, ***p<0.001, non-significant differences are not labeled. **I** Transmission electron microscope images of 14 days old mammospheres formed by parental, PMCA4-specific shRNA (sh-PMCA4), GFP-PMCA4b and GFP-PMCA4b^LA^-expressing MCF-7 cell lines cultured in Matrigel. Insets show magnified areas indicated by the dashed black squares.

**Figure 8.**
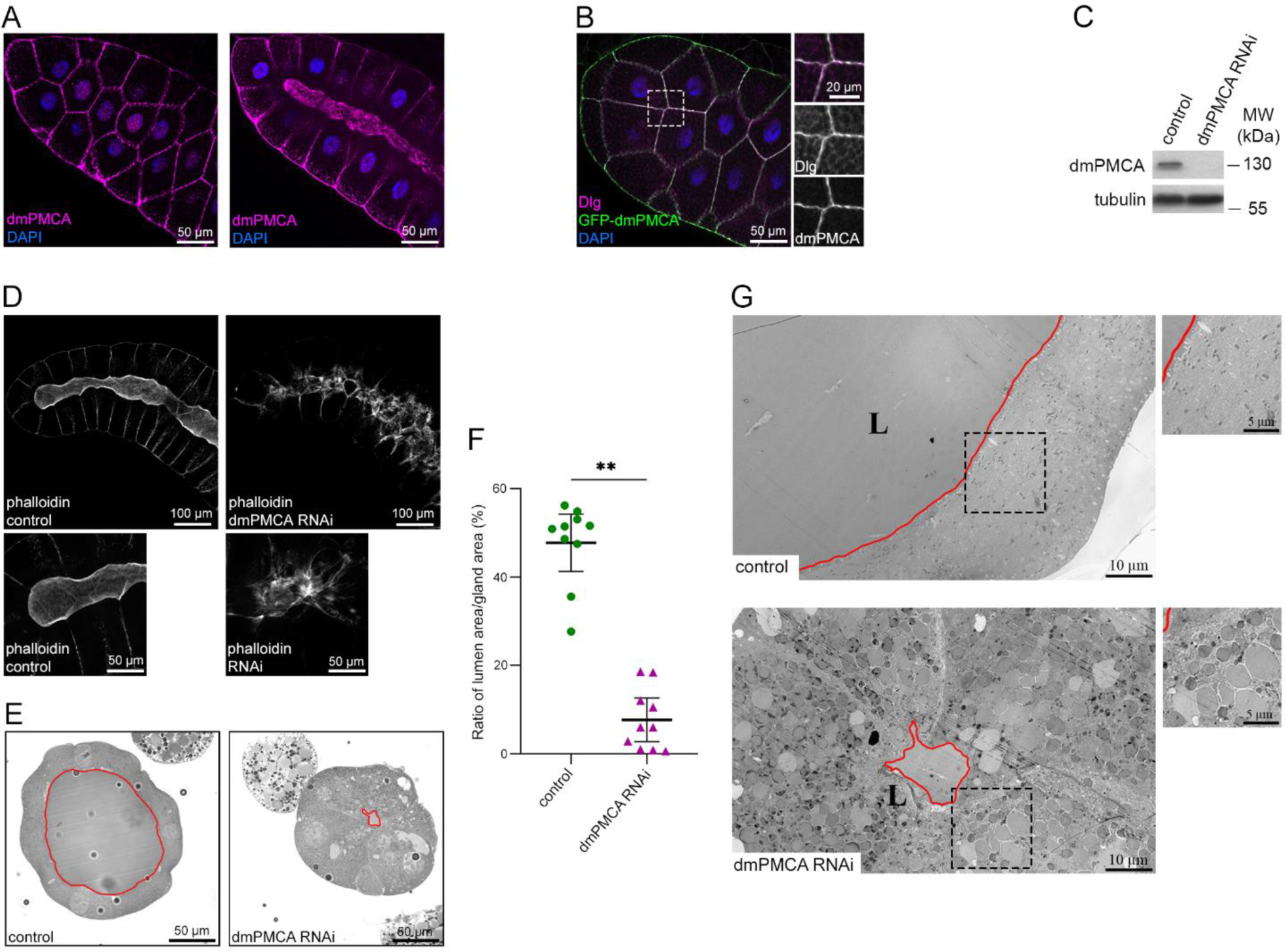
*Drosophila melanogaster* (dm)PMCA regulates lumen size and morphology in larval salivary gland. **A** DmPMCA immunostaining of the salivary gland of a 3^rd^ stage larvae. On the left: upper cell layer section; on the right: middle section. **B** Co-immunostaining of *Drosophila melanogaster* Dlg and (dm)PMCA of the salivary gland of a 3^rd^ stage larvae. **C** Western blot analysis of dmPMCA expression in control and ubiquitous *daughterless* promoter-driven dmPMCA-specific RNAi-expressing 3^rd^ stage *Drosophila melanogaster* larvae. **D** Phalloidin staining of control and salivary gland specific *forkhead* promoter-driven dmPMCA RNAi-expressing 3^rd^ stage larvae. Images in the lower row show distal parts of the lumen in the same salivary gland as above using higher magnification. **E** Toluidine blue-stained semi-thin cross-sections of salivary glands from 3^rd^ stage control and *forkhead* promoter-driven dmPMCA specific RNAi-expressing larvae. The lumen is outlined by red line. **F** Statistical comparisons of the lumen area ratios on semi-thin cross-sections. Graph displays means with 95% CI, n=10. Data were analyzed with Kruskal-Wallis and Dunn’s multiple comparisons tests, adjusted p values: ^ns^p≥0.05, **p<0.01. **G** Transmission electron microscope images from cross-sections of salivary gland of control and salivary gland specific *forkhead* promoter-driven dmPMCA RNAi-expressing 3^rd^ stage larvae. Magnified areas indicated by the dashed black rectangles are shown on the right. The lumen is outlined by red line. L: lumen.

To further study the role of PMCA in glandular development, we silenced dmPMCA using a dmPMCA-specific *UAS* promoter-driven RNA interference construct in the UAS-GAL4 system with *daughterless* promoter-driven Gal4 that effectively reduced dmPMCA expression in 3^rd^ stage larvae (Fig. 7C). Tissue-specific silencing of dmPMCA resulted in an aberrant F-actin pattern that normally borders the central lumen of the salivary gland (Fig. 7D). In addition, cross-sections of epoxy resin-embedded salivary gland samples showed a substantial reduction in central lumen size upon dmPMCA silencing (Fig. 7E, F). Closer inspection of the dmPMCA-deficient salivary gland cells by electron microscopy revealed a large secretory vesicle accumulation in the cytoplasm (Fig. 7G), suggesting secretion defect similar to that seen in the PMCA4-silenced MCF-7 mammospheres (Fig. 6I). These results highlight the evolutionarily conserved role of PMCA in the regulation of lumen morphology and merocrine secretion.

## Discussion

Altered histological differentiation in malignancies (including breast cancer) characterized by the loss of tissue architecture and loss of cell polarity is associated with poor patient survival (Enane et al., 2018). Normal mammary gland luminal epithelial cells exhibit apico-basal polarity, and this is progressively lost during tumorigenesis (Halaoui et al., 2017). In the current report, we show for the first time that PMCA4 is downregulated in tumor tissue samples collected from patients with luminal breast cancer when compared to healthy breast tissue, where PMCA4 expression is high. In addition, we also demonstrate that elevated PMCA4 expression is associated with longer relapse free survival in LUMA and LUMB1 breast cancer subtypes (Fig. 1). To study whether the severe loss of PMCA4b in luminal breast cancer cells can be related to the loss of cell polarity we either silenced or increased the level of PMCA4b in MCF-7 cells. Silencing of PMCA4b caused cytoplasmic localization of E-cadherin while it did not affect the expression of epithelial and mesenchymal markers or cell growth (Fig. 2), which are typical signs of partial EMT (Aiello et al., 2018). It is worth noting that low PMCA4 protein abundance found in a variety of tumor cells (including MCF-7) is associated with prolonged Ca^2+^ signaling (Hegedus et al., 2017b; Padanyi et al., 2016; Varga et al., 2014), and prolonged Ca^2+^ signaling has been implicated in partial EMT (Norgard et al., 2021). Here we show that the presence of PMCA4b in MCF-7 cells is necessary for the plasma membrane localization of E-cadherin and Dlg1 (Fig 2, 3), and that elevated levels of PMCA4b increase the number of polarized cells and induce polarized endosomal vesicular trafficking (Fig. 3, 4), suggesting that PMCA4b expression enhances cell polarization possibly through its interaction with the PDZ polarity protein Dlg1 (Fig. 9). Interestingly, loss of Dlg1 has also been implicated in cancer progression in a variety of tumors suggesting the importance of its proper expression and/or localization (Catterall et al., 2020; Fuja et al., 2004; Marziali et al., 2019). It is notable, that in contrast to our findings in gastric cancer cells PMCA4 silencing induced full EMT suggesting cancer tissue-specific function of PMCA4 (Wang et al., 2020).

**Figure 9.**
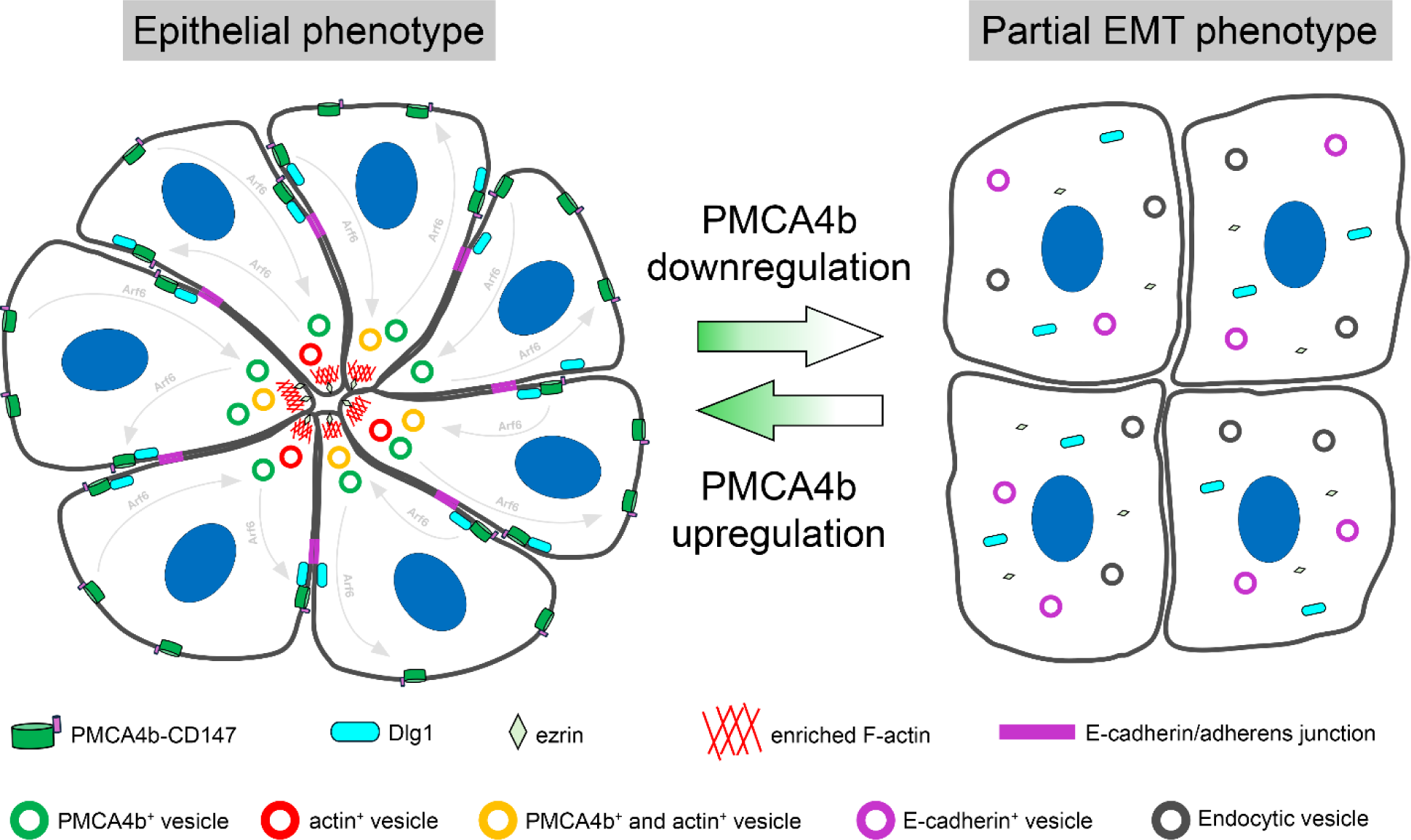
A model on the role of PMCA4b in luminal epithelial cell polarization and lumen formation. PMCA4b facilitates the lumen forming capability of luminal epithelial cells through contributing to the polarized distribution of ezrin, and the plasma membrane localization of E-cadherin and Dlg1. During (pre-)lumen formation dynamic vesicular trafficking toward the emerging lumen area includes trafficking of PMCA4b through its Arf6-mediated recycling to the basolateral plasma membrane compartment. Losing PMCA4b, epithelial cells lose their polarized phenotype and acquire partial EMT characterized by cytoplasmic localization of E-cadherin, Dlg1 and ezrin.

Loss of epithelial cell polarity is a hallmark of progression to DCIS, in which malignant cells lack the ability to form normal ductal structures, but they still express the epithelial marker E-cadherin (Chatterjee and McCaffrey, 2014). Here we show that PMCA4b over-expressing MCF-7 cells can create mammospheres with distinct central lumens surrounded by a single layer of cells, whereas PMCA4-silenced and PMCA4b^LA^-expressing MCF-7 cells formed microlumens typically found in DCIS (Fig. 7) (Chatterjee and McCaffrey, 2014). Higher capability of PMCA4b-expressing cells to form a central lumen is also supported by the significantly elevated proportions of pre-lumen-like structures in 2D cell cultures compared to the parental or PMCA4-silenced MCF-7 cells (Fig. 6). Silencing the single *Drosophila melanogaster pmca* gene resulted in larval salivary gland lumen morphology degeneration (Fig. 8), suggesting an evolutionarily conserved role of PMCA proteins in lumen formation of glandular organs *in vivo*. It has been demonstrated that secretory activity requires cytoplasmic Ca^2+^ oscillations (Ahuja et al., 2020; Mikoshiba et al., 2008; Pedersen et al., 2019), and local elevation of Ca^2+^ concentration is necessary during fusion of secretory granules with the plasma membrane (Xue et al., 2021). Our electron microscopy analysis of PMCA(4)-deficient MCF-7 mammospheres and Drosophila salivary glands revealed a pronounced secretion defect and abnormal accumulation of secretory vesicles both *in vitro* (Fig. 7D) and *in vivo* (Fig 8G). Revealing the exact PMCA function in secretion requires, however, further investigation.

Recently, we demonstrated that PMCA4b over-expression induced cell polarization and actin cytoskeleton remodeling of BRAF mutant melanoma cells. Moreover, a profound change in cell culture morphology and F-actin rearrangement was detected in MCF-7 cells in response to changes in PMCA4b protein abundance (Naffa et al., 2021). Actin network formation and contraction are prerequisites of lumen formation (Taniguchi et al., 2015) and several proteins involved in actin cytoskeleton dynamics are regulated by Ca^2+^ (Lehne and Bogdan, 2023). *In vitro* PMCA4 can directly interact with G-actin and short actin oligomers. Both interactions activate PMCA ATPase function while interaction with F-actin inhibits its activity (Dalghi et al., 2013; Vanagas et al., 2013). PMCA4b can also affect F-actin dynamics indirectly by reducing cytoplasmic Ca^2+^ concentration that may prevent cortical F-actin degradation in response to an uncontrolled rise in Ca^2+^ (Yoneda et al., 2000). It has been demonstrated that an enrichment of PMCA4 at the cell front was essential for the back to front Ca^2+^ gradient in controlling F-actin dynamics at the lamellipodia during endothelial cell migration, however, the exact mechanisms for directing PMCA4 to the cell front has not been elucidated (Tsai et al., 2014).

Our results highlight the importance of proper intracellular trafficking of PMCA4b in cell polarization and subsequent lumen formation and suggest that the small GTPase Arf6 may play a role in this process. We show that PMCA4b colocalizes both with the wild type and a constitutively active form of Arf6, and that Arf6 inhibition interferes with PMCA4b recycling. Previous studies indicate that Arf6 plays a role in actin cytoskeleton remodeling and lumen formation in epithelial and endothelial cells (Francis et al., 2023; Milanini et al., 2018; Monteleon et al., 2012; Tushir et al., 2010; Zangari et al., 2014) that is in good agreement with our current observation demonstrating that Arf6 inhibition prevent PMCA4b-induced pre-lumen formation in 2D cell cultures (Fig. 6E, F).

Our model shown in Figure 9 suggests that elevated PMCA4b levels promote the polarization of luminal epithelial cells with proper plasma membrane localization of E-cadherin, Dlg1 and ezrin. PMCA4b is involved in dynamic endocytic vesicle trafficking towards the central pre-lumen area where it may participate in central actin stabilization at the early steps of lumen formation (Fig. 6B). This is followed by its recycling to the basolateral plasma membrane of epithelial cells in fully developed luminal structures (Fig. 7A). During these processes PMCA4b trafficking is regulated by Arf6, in which the recently described PDZ protein complex PLEKHA7–PDZD11 may play a role (Sluysmans et al., 2022).

Loss of cell polarity and cell-cell contacts is a hallmark of epithelial malignancy (Chatterjee and McCaffrey, 2014; Peglion and Etienne-Manneville, 2024). Taken together with our observations on clinical outcome data (Fig. 1) this suggests that elevated levels of PMCA4b may be associated with a lower risk of cancer progression at early stages of luminal breast cancer. It is important to note, however, that different cancer types have different molecular backgrounds and hence, PMCA4b may play distinct roles. For instance, in contrast to our findings in melanoma cells (Hegedus et al., 2017a), PMCA4b knockdown reduced the migratory activity of pancreatic ductal adenocarcinoma MIA PaCa-2 cells and increased their sensitivity to apoptosis (Sritangos et al., 2020). It is worth mentioning that in pancreatic ductal cells PMCA4 localizes to the apical plasma membrane where it physically interacts with CFTR protein supporting its cell-type specific function (Madacsy et al., 2022). We also show here that whereas PMCA4 abundance is associated with longer relapse-free patient survival in luminal LUMA (grade 2) and LUMB1 breast cancer subtypes, the opposite is observed for patients with HER^+^ positive LUMB2 breast cancer (Fig. 1D). Thus, PMCA4 may have cancer type-specific functions in various neoplastic processes, emphasizing the importance of precise histopathological and molecular characterization of individual tumor samples for optimal treatment choices.

In summary, our findings suggest that loss of PMCA4 is implicated in luminal type breast carcinomas, and that elevated PMCA4 levels are associated with longer relapse-free patient survival at the early onset of LUMA and LUMB1 breast cancer cases. Our results may indicate that PMCA4b is a novel polarity protein and point towards its potential evolutionarily conserved function in the formation of luminal structures in exocrine glandular organs.

## Material and Methods

### Tissue samples

109 formalin fixed paraffin embedded HR^+^ (LUMA n=39; LUMB1 n=50; LUMB2 n=20 study cohort) and 4 normal breast tissue samples obtained after breast reduction surgery were selected (Supplementary Table 1). The cases were diagnosed at the Department of Pathology, Forensic and Insurance Medicine, Semmelweis University, Hungary between 2000 and 2010. Clinicopathological data of the patients were obtained from the files of the Semmelweis University, 2^nd^ Dept. of Pathology and from the Semmelweis University Health Care Database with the permission of the Hungarian Medical Research Council (ETT-TUKEB 14383/2017). Surrogate breast carcinoma subtype was defined based on four (estrogen receptor (ER), progesterone receptor (PgR), Ki67 index (marker of proliferation) and human epidermal growth factor receptor-2 (HER2)) immunohistochemical markers and according to the 2013 St. Gallen Consensus Conference recommendations (Goldhirsch et al., 2013).

Luminal A (LUMA) tumors are defined as ER and PgR positive, HER2 negative, Ki-67 “low” (Ki-67 < 20%) tumors, Luminal B-HER2 negative (LUMB1) tumors as ER positive, HER2 negative and Ki-67 “high” (≥20%) and/or PgR “negative or low” (PgR cut-point = 20%) and Luminal B-HER2 positive (LUMB2) as ER positive and HER2 overexpressed or amplified.

### Immunohistochemistry to evaluate PMCA4 expression in breast tissue samples

Immunohistochemistry (IHC) staining on all human tissue samples was performed and evaluated at the Department of Pathology, Forensic and Insurance Medicine, Semmelweis University. Cut-off values for estrogen receptor (ER) and progesterone receptor (PgR) positivity were 1% of tumor cells with nuclear staining and defined as positive or negative. Human erb-b2 receptor tyrosine kinase 2 (HER2) status was determined either as protein overexpression or HER2 gene amplification detected by fluorescence in situ hybridization (FISH). Data were analyzed by Kruskal-Wallis and Mann-Whitney tests for comparison of quantitative variables by the statistical software SPSS V.25 (IBM Corp.).

PMCA4 IHC was performed with an automated immunostainer system (Ventana Benchmark Ultra, Roche Diagnostics) according to the manufacturer’s instructions. Antigen retrieval was performed with ULTRA CC1 antigen retrieval solution for 90 min at 95°C. The primary anti-PMCA4 JA9 mouse monoclonal antibody (P1494, Merck) was used at a 1:200 dilution at 42°C for 32 min. For antibody visualization, the UltraView DAB Detection kit (Ventana) was applied. PMCA4 stained slides were scanned with slide-scanner (PANNORAMIC® 1000 DX, 3DHISTECH Ltd., Hungary). PMCA4 IHC reaction was semi-quantitatively evaluated on digitized slides: 1) IHC score was defined as the percentage of positive epithelial cells counted on average in 3–5 high-power fields, 2) membrane positivity with or without cytoplasmic reaction was assessed. PMCA4 IHC scoring was performed on digitized slides by using the slide-viewing software CaseViewer (CaseViewer 2.3.0.99276, 3DHISTECH Ltd., Hungary).

### Reagents and treatments

Phalloidin-TRITC (P1951, Sigma-Aldrich) and NAV2729 (SML2238, Sigma-Aldrich) were dissolved in dimethyl sulfoxide (DMSO) and stored at −20°C at a concentration of 80 µM and a 4.5 mM, respectively. Phalloidin-TRITC was used for F-actin labeling on confluent 2D cultures and 3D mammospheres of MCF-7 cells at a concentration of 80 nM. In the experiments studying the role of Arf6, MCF-7 cells were treated with 6.75 µM NAV2729 for 24 or 48 hours. To label nuclei 2-(4-amidinophenyl)-1H-indole-6-carboxamidine (DAPI) was used at a concentration of 0.5 µM. For *Drosophila melanogaster* salivary gland staining Rhodamine Phalloidin (ab235138, Abcam) was used at a 1000x dilution in PBS according to the manufacturer’s instructions.

### Cell culture

MCF-7 (ATCC HTB-22) and HEK-293 cell lines were purchased from the American Type Culture Collection (ATCC). Cells were cultured in Dulbecco’s modified Eagle’s medium (DMEM) (DMEM-HXA, Capricorn Scientific) supplemented with 10% fetal bovine serum (FBS; ECS0180L, Euroclone), 2mM L-glutamine (STA-B, Capricorn Scientific), 100 U/ml Penicillin and 100 µg/ml streptomycin (PS-B, Capricorn Scientific) in a humidified 5% CO_2_ incubator at 37 °C. In order to maintain the stable expression of PMCA4b transgenes and of the shRNA construct, 100 ng/ml puromycin dihydrochloride (sc-108071, Santa Cruz Biotechnology) was added to the culture media.

### Generation of stable cell lines

PMCA4-specific shRNA MCF-7 cells were generated by using the PMCA4-specific shRNA plasmid (sc-42602-SH, Santa Cruz Biotechnology, Santa Cruz, CA, USA), as described previously (Naffa *et al*., 2021). GFP-PMCA4b and GFP-PMCA4b^LA^-expressing MCF-7 and GFP-PMCA4b-expressing HEK-293 cell lines were generated by stable transfection of the cells with *SB-CAG-GFP-PMCA4b-CAG-puromycin* or *SB-CAG-GFP-PMCA4b-LA-CAG-puromycin* constructs that contain a Sleeping Beauty transposon system, using the protocol described previously (Hegedus et al., 2017a; Naffa et al., 2021).

### Transgenic Drosophila melanogaster lines

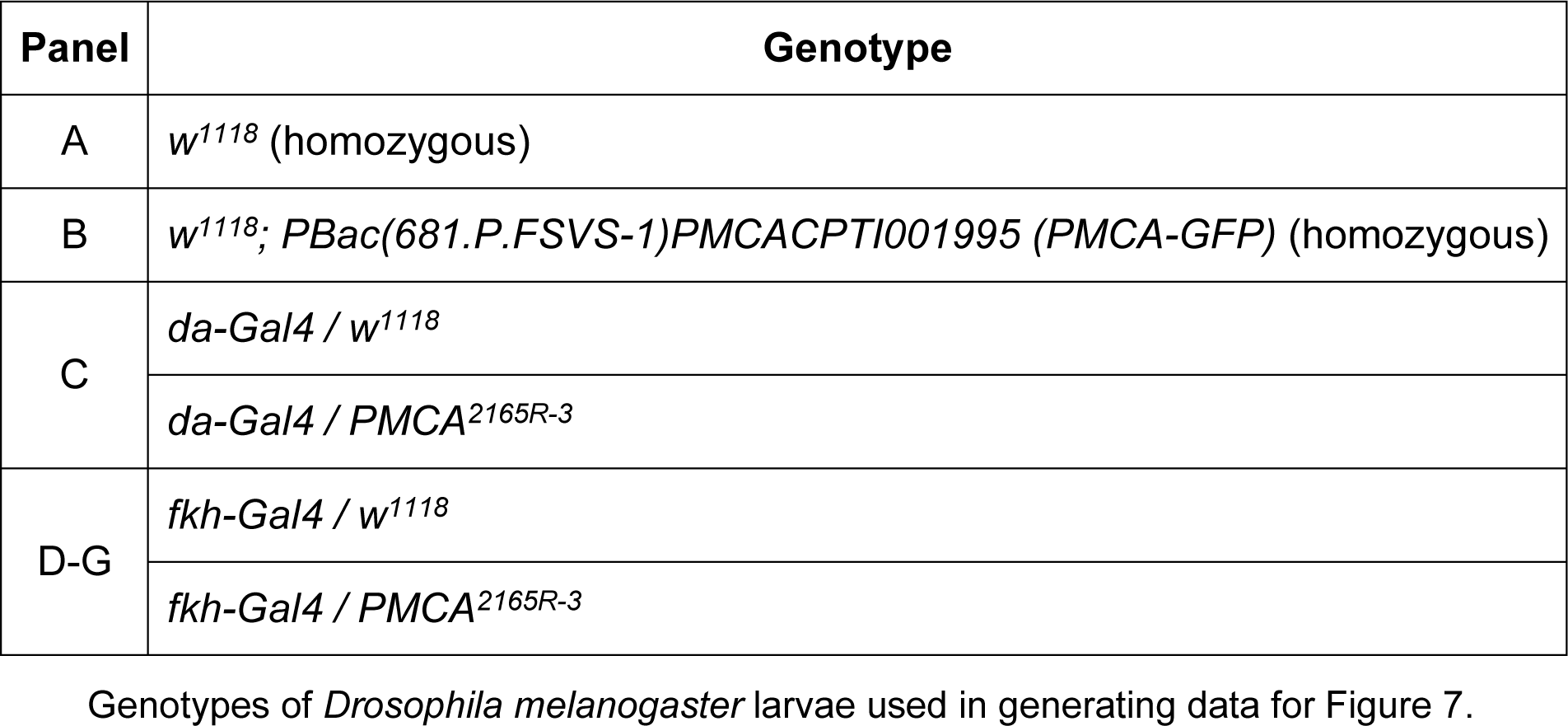

Flies were raised at 25°C on standard cornmeal, yeast and agar containing medium. The *w^1118^* strain (FlyBase ID: FBst0003605) was used as control and *da-Gal4* fly line (exact genotype: *w*;P(UAS-da.G)52.2*, FlyBase ID: FBst0051669) were obtained from the Bloomington Drosophila Stock Center (BDSC), the *PMCA^2165R-3^ RNAi* line (stock ID: 2165R-3) was obtained from the Fly Stocks of National Institute of Genetics (Nig-Fly) and the PMCA-GFP line was obtained from Kyoto Drosophila Stock Center (exact genotype: *w1118; PBac(681.P.FSVS-1)PMCACPTI001995*, DGRC number: 115256). The *fkh-Gal4* fly line was provided by Eric H. Baehrecke (Berry and Baehrecke, 2007) (University of Massachusetts Medical School, Worcester, MA).

### Immunofluorescence

#### 1. Immunocytochemistry of MCF-7 cells

Cells were seeded in removable 8-well chambers (80841, Ibidi) and 96 hours later rinsed with phosphate buffered saline (PBS) at 37°C, and fixed with 4% PFA diluted in PBS for 10 min. After rinsing twice and washing twice with PBS for 5 min, cells were permeabilized for 10 min. with 0.2% Triton X-100 diluted in PBS. After rinsing twice and washing twice with PBS, the cells were treated with Image-iT™ FX Signal Enhancer (I36933, Invitrogen) for 30 min to reduce non-specific binding of fluorophores. After rinsing twice with PBS, the samples were incubated in blocking solution containing 2 mg/ml bovine serum albumin (A7030, Sigma-Aldrich), 1% gelatin from cold water fish skin (G7765, Sigma-Aldrich), 0.1% Triton X-100 (X100PC, Sigma-Aldrich) and 5% goat serum (G9023, Sigma-Aldrich) diluted in PBS, for 1 hour at room temperature (RT). Samples were then incubated in the following primary antibodies diluted in blocking solution: anti-E-cadherin (24E10, CST 3195, Cell Signaling, dilution 1:200), anti-ezrin (E1281, Sigma-Aldrich, dilution 1:200), anti-Dlg1/SAP97 (sc-25661, Santa Cruz Biotechnology, dilution 1:100), anti-CD147/basigin (10186-R125, Sino Biological, dilution 1:200), or anti-HA (H3663, Sigma-Aldrich, dilution 1:500). After rinsing twice and washing twice for 5 min with PBS, the Alexa Fluor 594-conjugated goat anti-rabbit and anti-mouse IgG secondary antibodies (A-11012 and A-11005, Invitrogen, dilution 1:500) were added. Samples were rinsed twice and washed twice with PBS for 5 min and the nucleus was labeled with DAPI (details in section *Reagents and treatments*). The silicon part of the chamber was then removed, and the samples were mounted with Ibidi Mounting Medium (50001, Ibidi).

#### 2. Phalloidin staining of MCF-7 cells

Cells were seeded, grown, fixed, permeabilized and mounted as described in the previous section. Phalloidin-TRITC (P1951, Sigma-Aldrich) was added to the cells after permeabilization and incubated for 1 hour.

#### 3. IHC staining of Drosophila larval salivary gland

Salivary glands were dissected from late L3 stage Drosophila larvae in PBS and pre-permeabilized for 20 sec in 0.05% Triton X-100 diluted in PBS. Salivary glands were then fixed for 40 min at room temperature in 4% PFA diluted in PBS, rinsed twice and washed twice with 0.1% Triton X-100 diluted in PBS (PBST) for 15 min and incubated for 40 min at room temperature (RT) in blocking solution that contained 10% fetal bovine serum and 0.1% Triton X-100 diluted in PBS; then salivary glands were incubated for 24 hours at 4°C with the following primary antibodies: anti-*pan*-PMCA (5F10, MABN1802, Sigma-Aldrich, dilution 1:250), anti-Dlg (4F3, Developmental Studies Hybridoma Bank, dilution 1:100), anti-GFP (A10262, Invitrogen, dilution 1:1000) diluted in blocking solution. After rinsing twice and washing twice the samples with PBST for 15 min, they were incubated with the secondary antibodies diluted in blocking solution: Alexa Fluor 488-conjugated goat anti-chicken IgY (A-11039, Invitrogen, dilution 1:1000) and Alexa Fluor 594-conjugated goat anti-mouse IgG (A-11020, Invitrogen, dilution 1:1000). After rinsing twice and washing twice the samples with PBST for 15 min, nuclei were labeled with DAPI (as in section *Reagents and treatments*) and samples were mounted with Vectashield (H-1000-10, Vector Laboratories).

#### 4. Phalloidin staining of Drosophila larval salivary glands

Salivary glands were dissected, permeabilized and fixed the same way as described in the previous section. For F-actin labeling, the samples were incubated with Rhodamine Phalloidin (details in section *Reagents and treatments*) for 1 hour, then washed, nuclei labeled with DAPI and mounted the same way as described for IHC.

### Imaging

Fluorescent images were obtained with an Axio Imager M2 microscope (ZEISS) with an ApoTome2 grid confocal unit (ZEISS) using a Plan-Apochromat 63×/1.4 Oil (ZEISS) objective for human cells, a Plan-Apochromat 40×/0.95 Air (ZEISS) objective for semi-thin sections of Drosophila salivary glands, a Plan-Neofluar 20x/0.5 Air objective for mammospheres and Drosophila salivary glands with an Orca Flash 4.0 LT sCMOS camera (Hamamatsu Photonics) using the Efficient Navigation 2 software (ZEN 2) (ZEISS). Images in Figure 6A were obtained with LSM 710, AxioObserver confocal laser-scanning microscope (ZEISS) using a Plan-Apochromat 63×/1.4 Oil DIC M27 objective (ZEISS). Images in Figure 6D and Supplementary Figure 5C were obtained with a Nikon Eclipse Ti2 confocal microscope with a Plan-Apochromat 60×/1.4 Oil objective.

Time-lapse video imaging was performed on an inverted Nikon Eclipse Ti2 microscope with 488 nm and 638 nm laser lines for the Yokogawa CSU-W1 spinning disc scan head equipped with two back illuminated Photometrics Prime BSI scientific CMOS cameras for detection. Green and red fluorescence were recorded using 525/50 nm and 641/75 nm emission filters. For the recording, we used a NIS Elements AR 5.4 software with a dimension of 1024×1024 pixels and bit depth of 16 bits. For acquisition, we used a 40x ApoLambda/NA1.15 water-dipping objective. The videos were captured for 25 minutes. Final videos were created by the Image J software (Rueden *et al*, 2017).

### Growing and staining mammospheres

5000 cells from the parental, GFP-PMCA4b, GFP-PMCA4b^LA^ and PMCA4-specific shRNA-expressing MCF-7 lines were mixed with cold Matrigel (Corning Matrigel, DLW356231, Sigma-Aldrich) and 15 µl of cell-Matrigel mixtures were added to removable 8-well chambers (80841, Ibidi) and incubated at 37 °C for Matrigel polymerization. After 15 min, FBS supplemented DMEM was added to the wells, and cells were grown for 10 days until they formed mature mammospheres. For F-actin staining the samples were washed with PBS at 37 °C, fixed for 40 min with 4% PFA diluted in PBS, permeabilized for 20 min with 0,2% Triton X-100 in PBS, and after washing TRITC-Phalloidin was added for F-actin staining (see further details in section *Reagents and treatments*). Samples were mounted with Ibidi Mounting Medium.

### Transfection

GFP-PMCA4b-expressing MCF-7 (5×10^4^) and HEK293 (7×10^4^) cells were seeded in removable 8-well chambers (80841, Ibidi). Next day cells were transiently transfected with the following plasmids: *pcDNA3/hArf6(WT)-mCherry* (Addgene plasmid #79422) and *pcDNA3/hArf6(Q67L)-HA* (Addgene plasmid # 79425) (the plasmids were a gift from Kazuhisa Nakayama (Makyio *et al*, 2012)) using the FuGENE HD Transfection Reagent (E2311, Promega) according to the manufacturer’s instructions. 24 hours after transfection, cells were fixed, permeabilized, stained with DAPI and mounted. In the case of Arf6^Q67L^ the cells were fixed and labeled with anti-HA antibody on the next day (details are the same as described in the section *Immunofluorescence*).

### WGA uptake assay

5×10^4^ cells from the parental, GFP-PMCA4b, GFP-PMCA4b^LA^ and PMCA4 specific shRNA-expressing MCF-7 lines were seeded in removable 8-well chambers (80841, Ibidi) and were grown for 5 days in FBS-supplemented DMEM. Cells were rinsed with cold (4°C) HBSS (Hanks′ Balanced Salt solution; H15-009, PAA Laboratories) twice and incubated with wheat germ agglutinin (WGA; W21404, Life Technologies) at a concentration of 5 µg/ml in HBSS for 10 min at room temperature. Unbound WGA was removed by rinsing the cells twice with PBS. The cells were then fixed, permeabilized and stained with DAPI, as described in the section *Immunofluorescence*.

### Sulforhodamine B (SRB) Assay

MCF-7 cells at densities indicated in Supplementary Figure 2B were seeded in a 96-well plate and cultured for 96 hours. Then cells were fixed with 10% trichloroacetic acid at 4 °C for 1 hour and washed with distilled water. After air dried overnight, 0.4% SRB was added to the wells and incubated for 15 min at room temperature. Excess stain was removed by washing with 1% acetic acid solution, and the bound SRB dye was solubilized in 10 mM Tris base solution for 10 min at RT with agitation. Optical density was measured at 570 nm using a microplate reader (EL800, BioTek Instruments, Winooski, VT, USA).

### Western blot analysis

The total proteins of MCF-7 and A375 cells were extracted by using the protocol described previously (Hegedus et al., 2017b). *Drosophila melanogaster* third-instar larvae were collected, homogenized in Laemmli buffer, and centrifuged to separate the protein containing supernatant. Equal amounts of proteins were loaded on 10% polyacrylamide gel, electrophoresed and electroblotted onto polyvinylidene difluoride (PVDF) membrane. Primary antibodies used for immunostaining were: anti-PMCA4 (JA9, P1494, Sigma-Aldrich, dilution 1:1000), anti-*pan*-PMCA (5F10, MABN1802, Sigma-Aldrich, dilution 1:2500), anti-E-cadherin (CST 3195, Cell Signaling, dilution 1:1000), anti-N-cadherin (sc-8424, Santa Cruz Biotechnology, dilution 1:500), anti-vimentin (M7020, Dako Products, dilution 1:1000), anti-β-actin (A1978, Sigma-Aldrich, dilution 1:2000), anti-Dlg1/SAP97 (sc-25661, Santa Cruz Biotechnology, dilution 1:1000), anti-CD147/basigin (10186-R125, Sino Biological, dilution 1:1000), anti-β-tubulin (ab6046, Abcam, dilution 1:1000). Horseradish peroxidase-conjugated secondary antibodies were used (715-035-151 and 711-035-152, Jackson ImmunoResearch, dilution 1:10,000) for detection with Pierce ECL Western Blotting Substrate (Thermo Scientific) and luminography on CL-XPosure Film (Thermo Scientific) or Amersham Hyperfilm ECL (GE Healthcare) films.

### Transmission electron microscopy (TEM)

MCF-7 cells were grown in Matrigel as described above for 14 days to create mammospheres. After the medium was removed, mammospheres were washed with FBS-free DMEM and fixed in a 3.2% methanol free, ultrapure formaldehyde, 0.25% glutaraldehyde, 0.029 M sucrose, 0.002 M CaCl_2_ and 0.1 M sodium cacodylate containing fixing solution for 24 hours at 4 °C. Samples were then washed for 2×5 min with 0.1 M sodium cacodylate, post-fixed in 1% ferrocyanide-reduced OsO_4_ diluted in 0.1 M sodium cacodylate, pH 7.4, at 4 °C for 1 hour and washed again for 2×5 min with 0.1 M sodium cacodylate. For contrasting, samples were washed for 2×5 min with ultrapure distilled water and incubated in 1% uranyl-acetate diluted in distilled water for 30 min. Then samples were dehydrated by using a graded series of ethanol and were embedded in Spurr low viscosity epoxy resin (EM0300, Sigma Aldrich). 70-nm ultrathin sections were made with an Om U3 microtome (Reichert Austria), collected on copper grids (AGG214, Agar Scientific) and stained in Reynold’s lead citrate. For examination we used a transmission electron microscope (JEM-1011; JEOL, Tokyo, Japan) equipped with a digital camera (Morada; Olympus) using iTEM software (Olympus).

### Production of semi-thin cross-sections of *Drosophila melanogaster* larvae

Salivary glands were dissected from late L3 stage Drosophila larvae then fixed in a 3.2% methanol free, ultrapure formaldehyde, 0.25% glutaraldehyde, 0.029 M sucrose, 0.002 M CaCl_2_ and 0.1 M sodium cacodylate containing fixing solution for 24 hours at 4 °C. Samples were than immobilized in 1.5% agar dissolved in distilled water (this helps to keep orientation and integrity of the samples), and embedded in Spurr resin the same way as described in section *Transmission electron microscopy*. 0.5-1 µm semi-thin sections were made with an Om U3 microtome (Reichert Austria) and stained with a 1% toluidine blue, 1% Azure II, 1% borax and 25% sucrose containing solution.

### Image analysis and statistics

IHC data were analyzed by Kruskal-Wallis and Mann-Whitney tests for comparison of quantitative variables by statistical software SPSS V.25 (IBM Corp.).To compare the prognostic impact of *ATP2B4* (PMCA4) and *ATP2B1* (PMCA1) expression in different breast carcinoma subtypes the publicly available database KM Plotter Online Tool (http://www.kmplot.com) was used. Only JetSet best probe set, and best cutoff values were selected, and Relapse Free Survival (RFS) plots were visualized by the Kaplan– Meier plotter using microarray data analysis (Lanczky and Gyorffy, 2021). Data for PMCA4 (NP_001675.3:S328) and PMCA1 (NP_001673.2:S1182) protein expression in normal and luminal breast cancer subtypes were derived from the Clinical Proteomic Tumor Analysis Consortium (CPTAC) and analyzed by using the UALCAN web portal (Chandrashekar et al., 2022) (https://ualcan.path.uab.edu/analysis-prot.html). Student’s t test p-values were considered significant below 0.05.

To evaluate the number of E-cadherin- and Dlg1-positive vesicles, the fluorescent images from original, unmodified single focal planes were analyzed with the Image J software (Rueden et al., 2017). The fluorescence signal was set by auto threshold, transformed into binary format and particles were counted with the Analyze Particles command, in which particle circularity was set between 0.5 and 1, and edges were excluded. Data were normalized to the total cell number determined by the nuclear dye DAPI image. Number of pre-lumens in 2D cell cultures and lumens in 3D cultures were quantified manually. To quantify the size of the lumens in Drosophila salivary glands, we measured the total area of the gland, as well as the lumen area in the cross-section images using ZEN 2 software, and their ratio was determined by dividing the lumen area with the total area of the gland. Data were imported into the GraphPad Prism 8.0 (GraphPad Software, Boston, Massachusetts USA, www.graphpad.com) software and tested for normality of data distribution using D’Agostino & Pearson and Shapiro-Wilk tests. When data showed normal distribution with both tests, we used one-way ANOVA and Tukey’s multiple comparisons tests; when data showed non-normal distribution, we used Kruskal-Wallis and Dunn’s multiple comparisons tests.

To define the ratio of polarized MCF-7 cells in a 2D cell cultures, the number of cells with either asymmetric or symmetric distribution of the apical marker ezrin was quantified manually in 16 fields of view for all cell lines. The total number of cells of each type was collected into contingency tables and data was analyzed by the Chi-square test.

To analyze planar distribution of WGA positive vesicles in MCF-7 cells we defined the center of nucleus with the Centroid command of the Image J software. The center of nucleus was then connected to the WGA-positive vesicles and the angle of the lines was defined by Image J. The obtained data were imported into an Excel table. Because Image J defines angles in a range of −180° to +180°, the data were converted into the range of 0° to 360°, and the angles were normalized to 180°. Then we calculated the mean values for each cell and determined their deviation from 180°. This deviation was added to the angles of each corresponding cell to get the final dataset. If the obtained value was higher than 360°, we subtracted 360 from it, if it was lower than 0° it was corrected by the addition of 360. Then we calculated the percentage of data in each bin for the whole datasets. We evaluated at least 15 cells of each cell type, and the data were copied to the Minitab (v.21) (https://www.minitab.com) followed by distribution analysis and histogram generation. Data were analyzed by the Levene’s probe using the “Two sample variance test” for each combination of the groups as illustrated by a box plot diagram.

For *Homo sapiens* PMCA4b and *Drosophila melanogaster* PMCA amino acid sequence comparison the P23634-6 and Q9V4C7 sequences were used from UniProt (UniProt, 2023). Alignment was depicted and analyzed by the Jalview 2.11.3.1 software (Waterhouse et al., 2009) using the “Percentage Identity” dynamic color scheme. Conservation values were automatically calculated in Jalview as described in the paper of Livingstone and Barton (Livingstone and Barton, 1993).

The number of evaluated images (n) and experiments are indicated in figure legends.

## Acknowledgements

The authors wish to thank Mónika Truszka for her help with electron microscopy, and Dr. John T. Penniston for his continuous support. This research was supported by the National Research, Development and Innovation Office under grant numbers NRDI K135811 (to AE), PD135447 (to TCs) and TKP2021-EGA-24 (to AE and AT). The New National Excellence Program of the Ministry for Innovation and Technology from the source of the National Research, Development and Innovation Office (ÚNKP-23-5-ELTE-603 to TCs) and the János Bolyai Research Scholarship of the Hungarian Academy of Sciences (BO/00023/21/8 to TCs).

---

**Video 1**. Time-lapse spinning disc confocal microscopy of GFP-PMCA4b-expressing MCF-7 cells after WGA uptake assay. Cells were incubated with WGA for 10 minutes, washed and transferred to a 37 °C incubator for 20 minutes. After that GFP-PMCA4b (green) and WGA (red) signals were captured for 25 minutes with spinning disc confocal microscope. Frame rate is 10 FPS.

**Video 2**. Time-lapse spinning disc confocal microscopy of PMCA4 specific sh-RNAi-expressing MCF-7 cells after WGA uptake assay. Cells were incubated with WGA for 10 minutes, washed and transferred to a 37 °C incubator for 20 minutes. After that WGA (red) signal were captured for 25 minutes with spinning disc confocal microscope. Frame rate is 10 FPS.

**Video 3**. Time-lapse spinning disc confocal microscopy of GFP-PMCA4b^LA^-expressing MCF-7 cells after WGA uptake assay. Cells were incubated with WGA for 10 minutes, washed and transferred to a 37 °C incubator for 20 minutes. After that GFP-PMCA4b^LA^ (green) and WGA (red) signals were captured for 25 minutes with spinning disc confocal microscope. Frame rate is 10 FPS.

**Supplementary Figure 1.**
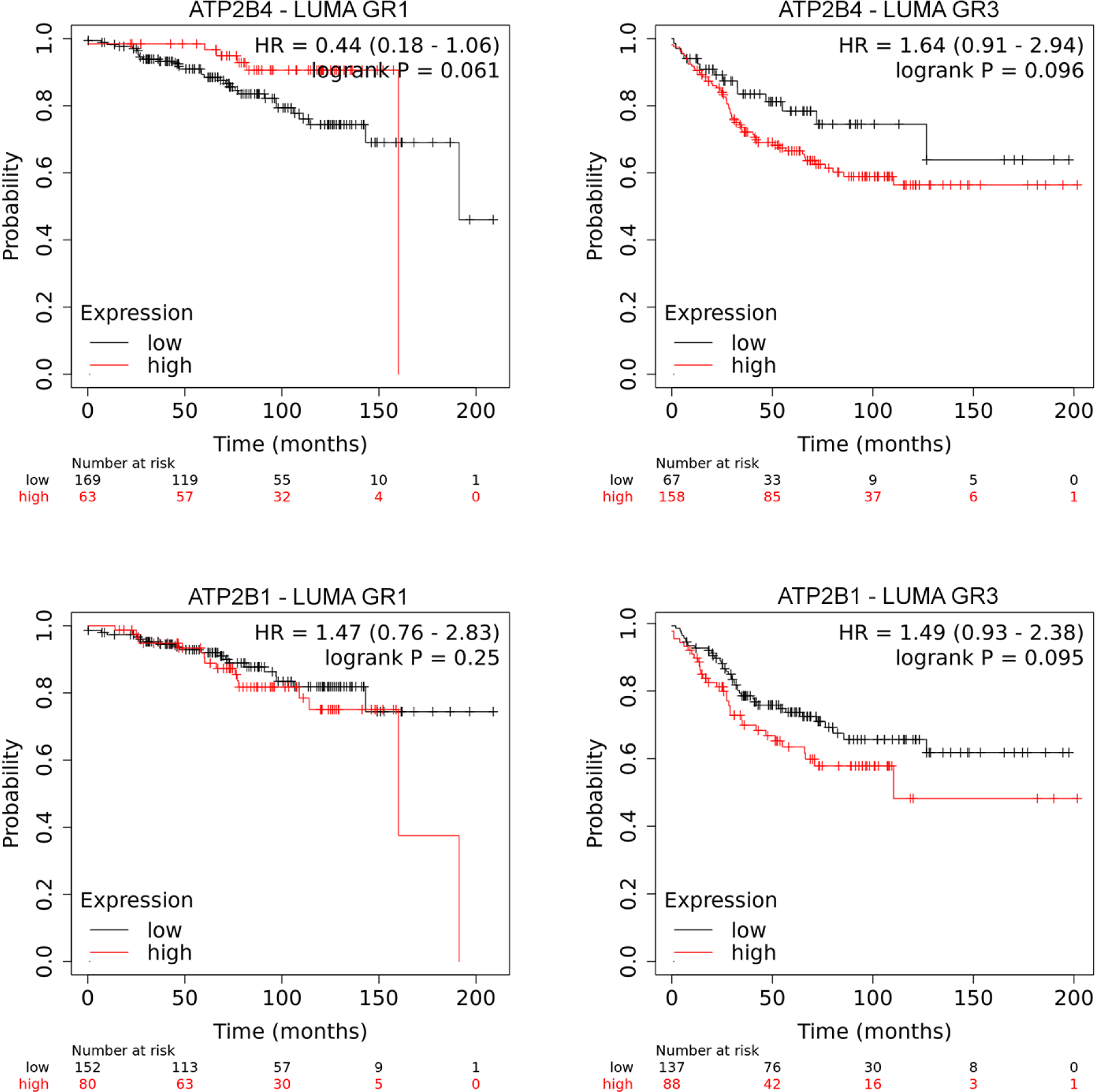
Kaplan–Meier relapse free survival analysis in breast cancer patients with LUMA subtype tumors with low and high *ATP2B4*/PMCA4 and *ATP2B1*/PMCA1 expression levels. GR1 and GR3 indicate grade 2 and 3. Data were collected by the online survival analysis tool (http://www.kmplot.com) using microarray data analysis; p values and the sample sizes are indicated in the figure.

**Supplementary Figure 2.**
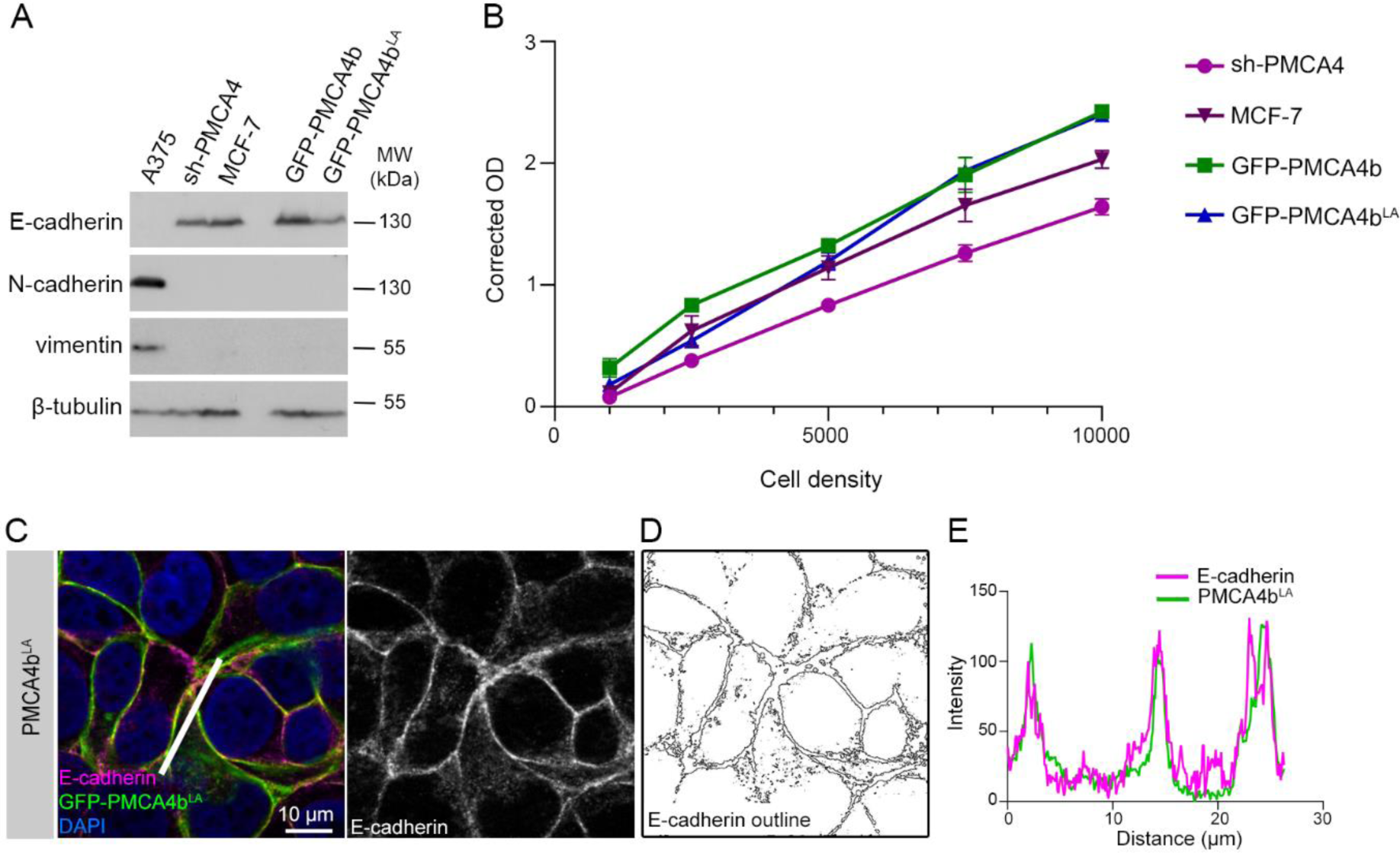
MCF-7 cells display epithelial characteristics. **A** Western blot analysis of epithelial (E-cadherin) and mesenchymal marker (N-cadherin, vimentin) proteins in mesenchymal-type A375 cells, and in epithelial-type PMCA4 specific shRNA-expressing, parental, GFP-PMCA4b and GFP-PMCA4b^LA^-expressing MCF-7 cell lines. A375 cells were used for N-cadherin and vimentin validation. **B** Cell density analysis of PMCA4 specific shRNA-expressing, parental, GFP-PMCA4b and MCF-7 cell lines by using sulforhodamine B assay. **C** E-cadherin immunostaining of GFP-PMCA4b^LA^-expressing MCF-7 cells. **D** Outline of the E-cadherin-positive cell compartments generated by the Image J software. **E** Fluorescence intensity profiles of E-cadherin and GFP-PMCA4b across the line shown in C.

**Supplementary Figure 3.**
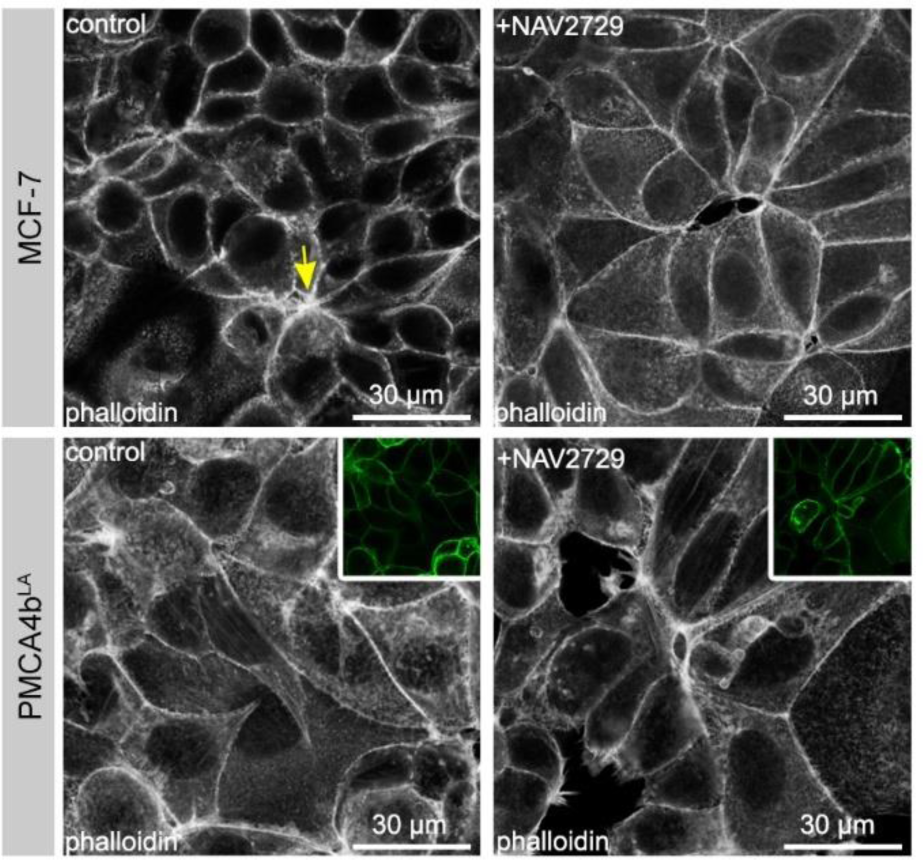
Effects of Arf6 inhibition on actin pattern and pre-lumen formation. Phalloidin staining of control and NAV2729-treated parental and GFP-PMCA4b^LA^-expressing MCF7-cells. Scaled-down insets show GFP-PMCA4b^LA^ signal. Arrow points to a pre-lumen.

**Supplementary Figure 4.**
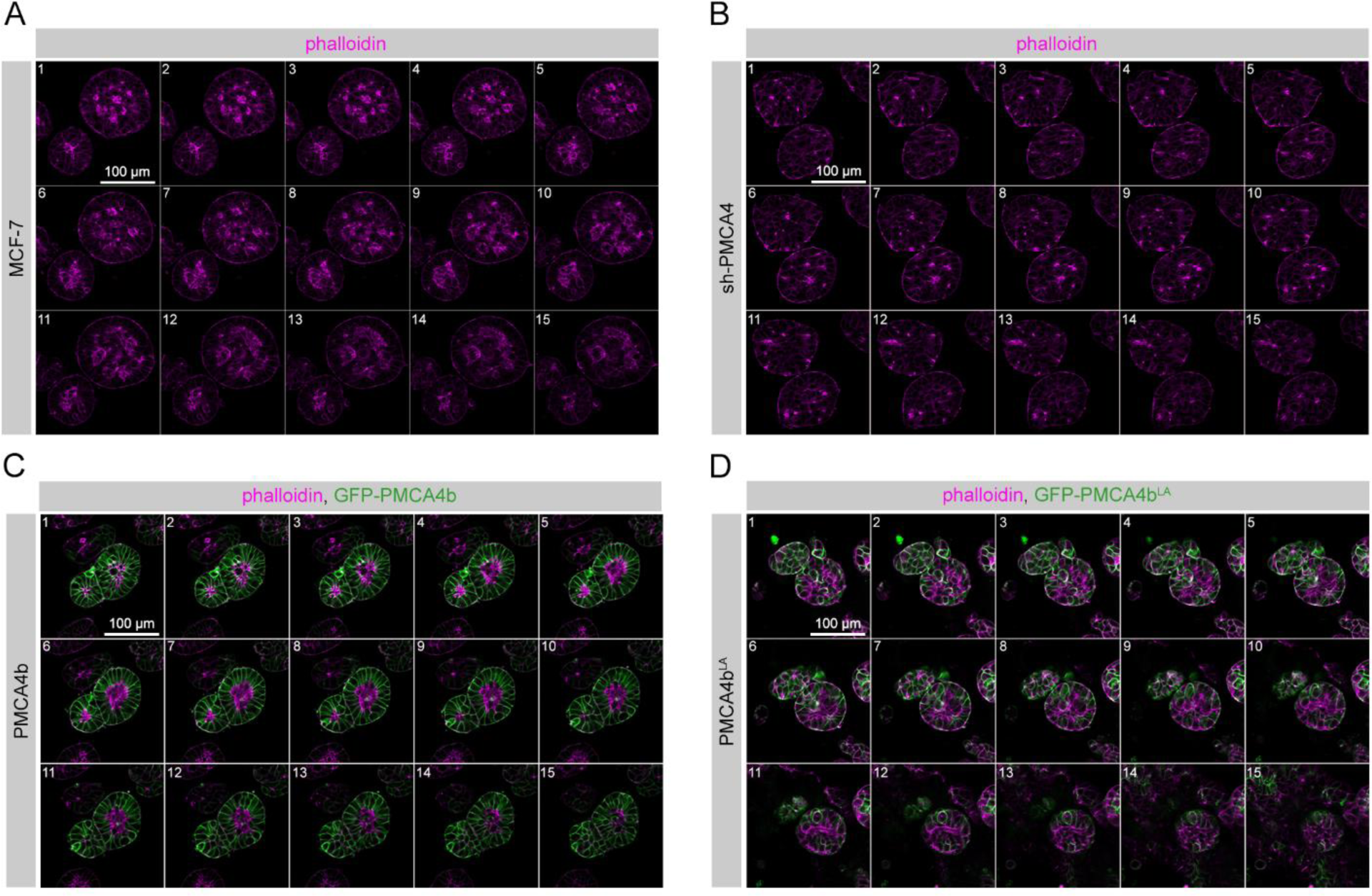
Lumen formation in MCF-7 mammospheres. **A-D** Z-stack images of parental, PMCA4 specific shRNA (sh-PMCA4), GFP-PMCA4b and GFP-PMCA4b^LA^-expressing MCF-7 cells grown in Matrigel for 10 days and stained with phalloidin. The range is 1.25 µm between slices.

**Supplementary Figure 5.**
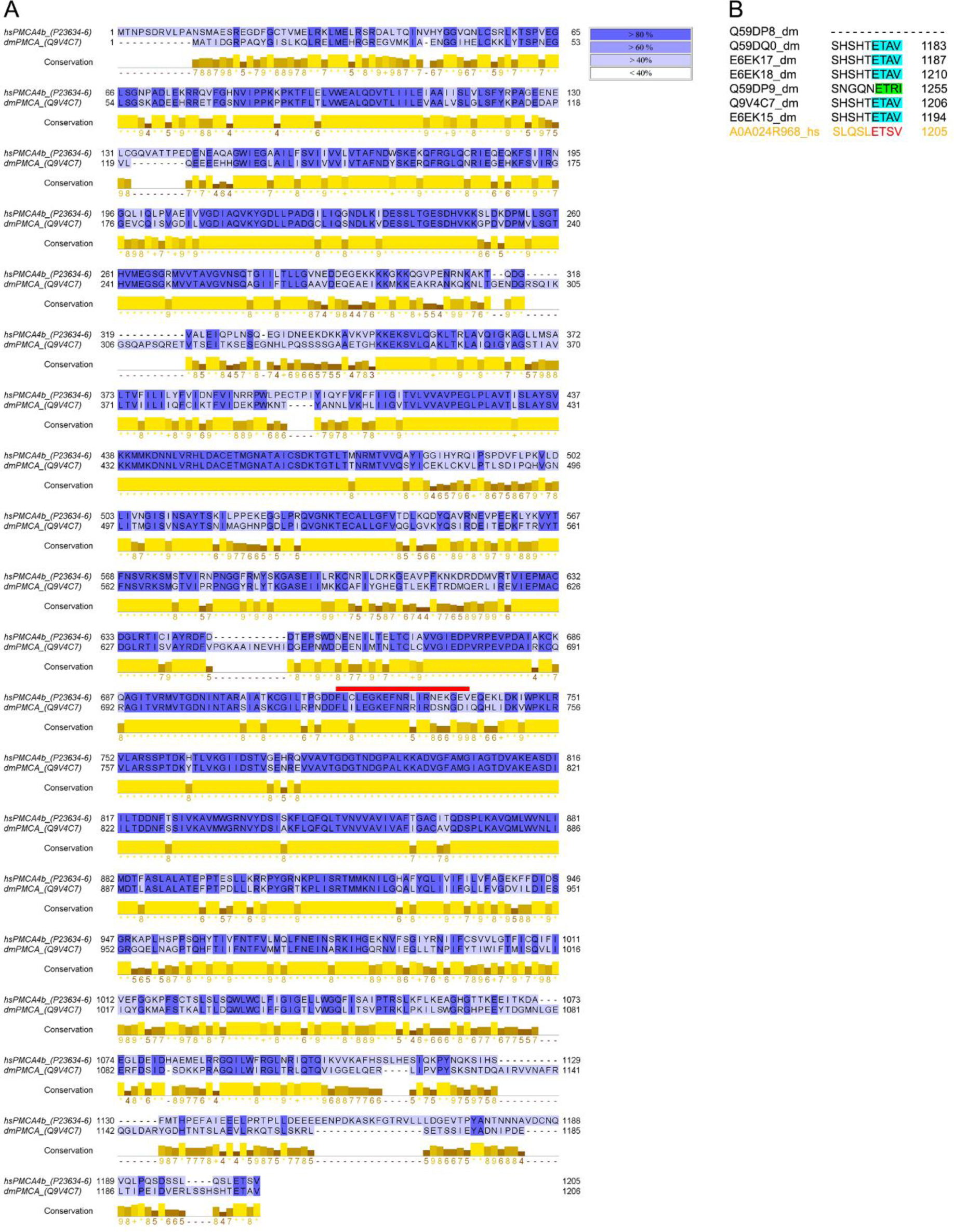
Amino acid sequence comparison of human PMCA4b and *Drosophila melanogaster* PMCA sequences. **A** Alignment of human PMCA4b and *Drosophila melanogaster* (dm)PMCA amino acid sequences using the Jalview software with the region recognized by the 5F10 antibody (located 719-738 in the human amino acid sequence) marked with red line. Purple colors indicate the rate of similarities between amino acids in the same position (ranges are indicated at right) with Percentage Identity Coloring in Jalview. Conservation row in yellow and brown shows conservation rate based on physico-chemical properties. Conserved columns are indicated by asterisks and columns with conserved changes are indicated by “+”. **B** Multiple sequence alignment of the C termini of dmPMCA isoforms (black) and human PMCA4b (yellow) by CLUSTAL O (1.2.4). Red color labels the PDZ-binding sequence of human PMCA4b, blue color labels Drosophila sequences with only 1 and green color labels 2 amino acid difference from human sequence.

**Supplementary Table 1.** Clinicopathological characteristics of the breast cancer patients’ samples. ER (estrogen receptor), PgR (progesterone receptor), HER2 (human erb-b2 receptor tyrosine kinase 2) status, and Ki67 (marker of proliferation Ki67; low means below 5%) index were evaluated by immunohistochemistry. The following data were taken into consideration: age at diagnosis, histological grade, pathologic tumor size (pT), nodal involvement (pN), surrogate breast carcinoma subtypes. T1: the tumor in the breast is 20 millimeters (mm) or smaller in size at its widest area; T2: the tumor is larger than 20 mm but not larger than 50 mm. N0: no cancer was found in the lymph nodes; N1: metastatic lymph nodes. Non-parametric statistical analysis was performed using Kruskal-Wallis and Mann-Whitney tests for comparison of quantitative variables. Statistical significance was defined at p<0.05.

|  |  | PMCA4 |  |  |  |
| --- | --- | --- | --- | --- | --- |
|  |  | Low | High | Total | P values |
| **ER** | Positive | 83 | 26 | 109 | - |
| **PR** | Negative | 22 | 6 | 28 | 0.727 |
|  | Positive | 61 | 20 | 81 |  |
|  | Total | 83 | 26 | 109 |  |
| **Ki67** | Low | 56 | 13 | 69 | 0.107 |
|  | High | 27 | 13 | 40 |  |
|  | Total | 83 | 26 | 109 |  |
| **HER2+** | Negative | 71 | 17 | 88 | 0.048 |
|  | Positive | 12 | 8 | 20 |  |
|  | Total | 83 | 25 | 108 |  |
| **Subtypes** | LUMA | 30 | 9 | 39 | 0.142 |
|  | LUMB1 | 41 | 9 | 50 |  |
|  | LUMB2 | 12 | 8 | 20 |  |
|  | Total | 83 | 26 | 109 |  |
| **Age** | Mean | 57.54 | 61.27 |  | 0.186 |
|  | SD | 12.59 | 12 |  |  |
|  | Range | 27-80 | 44-83 |  |  |
|  | Total | 83 | 26 | 109 |  |
| **Grade** | 1 | 23 | 6 | 29 | 0.519 |
|  | 2 | 41 | 10 | 51 |  |
|  | 3 | 18 | 8 | 26 |  |
|  | Total | 82 | 24 | 106 |  |
| **pT** | 1 | 49 | 13 | 62 | 0.492 |
|  | 2 | 33 | 12 | 45 |  |
|  | Total | 82 | 25 | 107 |  |
| **pN** | 0 | 49 | 14 | 63 | 0.923 |
|  | 1 | 33 | 9 | 42 |  |
|  | Total | 82 | 23 | 105 |  |
| **Metastatic** | Yes | 13 | 3 | 16 | 0.621 |
|  | No | 68 | 22 | 90 |  |
|  | Total | 81 | 25 | 106 |  |

